# Effectiveness of active learning for ecology teaching: the perspective of students vs their grades

**DOI:** 10.1101/2020.04.02.021584

**Authors:** Carlos Frankl Sperber, Neucir Szinwelski, Frederico Fernandes Ferreira, Lucas Ferreira Paiva, Victor Mateus Prasniewski, Ana Flávia de Paula Teixeira, Bruno Cabral Costa, Renata Bernardes Faria Campos, Rita Márcia Andrade Vaz de Mello, Benjamin Wiggins

**Affiliations:** Departmento de Biologia Geral (Professor at the General Biology Department), Universidade Federal de Viçosa, Viçosa, MG, Brazil; Centro de Ciências Biológicas e da Saúde (Professor at the Health and Biological Sciences Center), Universidade Estadual do Oeste do Paraná, Cascavel, PR, Brazil; Programa de Pós-graduação em Ecologia (Student at the Graduation Program in Ecology), Universidade Federal de Viçosa, Viçosa, MG, Brazil; Graduação em Engenharia Elétrica (Undergraduate student in Electrical Engineering), Universidade Federal de Viçosa, Viçosa, MG, Brazil; Programa de Pós-graduação em Conservação e Manejo de Recursos Naturais (Graduation Program in Conservation and Managements of Natural Resources), Universidade Estadual do Oeste do Paraná, Cascavel, PR, Brazil; Programa de Pós-graduação em Ecologia e Conservaço da Biodiversidade (Student at the Graduation Programm in Ecology and Conservation of Biodiversity), Universidade Federal do Mato Grosso, Cuiabá, MT, Brazil; Graduação em Arquitetura e Urbanismo (Undergraduate student in Architecture and Urbanism), Universidade Federal de Viçosa, Viçosa, MG, Brazil; Programa de Pós-graduação em Educação (Student at the Graduation Program in Education), Departamento de Educação, Universidade Federal de Viçosa, Viçosa, MG, Brazil; Universidade Vale do Rio Doce (Professor), Governador Valadares, MG, Brazil; Departamento de Educação (Professor at the Education Department), Universidade Federal de Viçosa, Viçosa, MG, Brazil; Department of Biology (Professor), Education University of Washington, Washington DC, USA

**Keywords:** cultural capital, peer instruction, cooperative learning, Paulo Freire, reflexive teacher, critical thinking, inclusive education, quantitative hypothesis testing, transformational learning, scientific teaching, effective learning in large classes

## Abstract

We evaluated the effectiveness of active learning for ecology teaching by comparing the perspective of students to their grades in exams. We estimated the perspective of the students through anonymous survey; we used students’ exam grades to estimate their ecology learning, and their effort and performance in the active learning tasks through their grades and proportion of intermediate steps for each active learning task. Active learning involved teachers’ stimuli for students’ active involvement, extra-class group task, individual online writing assessments, redoing exam in pairs, and classroom writing group assessments. We also evaluated the impact, unto the effectiveness of active learning, of several student characteristics, such as sex, age, individual study effort, and previous basic knowledge. We found that self-evaluated learning increased linearly with teachers’ attempts to stimulate students’ active involvement (*P* = 0.0003), extra-class group task (*P* = 0.0003), and previous basic knowledge (*P* = 0.02), while students’ grades increased asymptotically with extra-class group task (*P <* 2^−16^), and increased linearly with online writing assessments (*P* = 9.3^−8^) and classroom-based writing group assessments (*P* = 0.03). Our results showed that students perceive most part of the effectiveness of active-learning tasks and of teachers’ efforts. We showed that active learning tasks are complementary, so we recommend that teachers in both college and high school should implement simultaneous active-learning tasks, that include extra-class work in group, individual and group writing assessments, and should stimulate students’ engagement through respectful and non-authoritarian behavior of the teacher. Our results also showed that previous basic knowledge also plays a central role in driving effective learning, evidencing the importance of students instruction outside college. The applied teaching methodology is cheap and feasible for large classes. In these times of rising intolerance, prejudice, dismiss of environmental issues and disregard of science itself, we need an effective, pluralistic, respectful, and student-centred education, that fosters critical thinking, tolerance and respect for differing points of view.

## Introduction

Ecology is integral to curricula in several undergraduate programs, ranging from biologically centered programs like biological sciences and applied biology subjects (e.g. agronomy), to programs such as civil and electrical engineering. Motivated by an urge to foster effective learning and critical thinking, in the year 2000 the ecology teachers ^1^ at the Federal University of Viçosa’s General Biology Department introduced weekly classroom writing assessments, to be answered in groups, as an active learning task designed to enhance learning effectiveness and student attention levels [1]. In recent years, we intensified active learning by implementing: (i) teachers’ attempts to stimulate students’ active involvement, (ii) extra-class group task, (iii) online writing assessments, redoing exam in pairs, besides maintaining (v) classroom writing group assessments.

### Educational challenges in a colonized country

Ever since the Portuguese royal family first arrived in Brazil, the country’s education system served as an elitist institution aimed at rich white men [2]. In 2003, for example, only 5% of university students had family income per capita that reached the level of one minimum wage (ca. US$ 69 per month, US$ 891 per year), while 71% of the university students had wages of over US$ 4450 per year, contrasting to 60% of the population between 18 and 24 years old, which had not concluded elementary school, earning up to half the minimum wage, that corresponds to US$ 350 per year [3]. Similar inequalities were also present regarding the ethnic proportion of young people entering university: 21% white vs. 5% non-white, contrasting to 44% of self-declared not-white people in these age interval, that had not concluded elementary school [3].

Since 2000, several changes have been implemented, aiming to broadening and democratizing access to higher education in Brazil, such as institutionalization of the quota system, which reserves seats for formerly excluded social groups, the Restructuring and Expansion of Federal Universities - Reuni, the Unified Selection System - SiSU, and, in the private network, the expansion of Student Funding - FIES and the University for All Program – Prouni [4]. These changes altered the profile of Brazilian university students, and increased university student places, especially at public universities [5], which are generally considered the best institutions in Brazil. This has enabled an increase in the proportion of formerly excluded social groups, such as Afro-descendants, among the students in public universities. In 2018, self-declared Afro-descendants reached, for the first time in Brazilian history, majority (50.3%) among university students, although this is still bellow their representation in the population (55.8 %) [4, 6].

The improved access to universities has, however, not equated to success at these institutions [7]. Over 50% of university entrants in 2010 abandoned their course within five years [8], this high drop-out rate indicating the serious difficulties students face in meeting the standard expected of them. Even worse than that, the quality of middle and high school education in Brazil is very low: Brazilian students scored lower than the Organisation for Economic Co-operation and Development (OECD) average in reading, mathematics and science [9]. For example, only 2% of Brazilian students performed at the highest levels of proficiency in reading, mathematics or science, and 43% of students scored below the minimum level of proficiency in all three subjects.

One of the strong predictors for the Brazilian students’ low performance is socio-economic status: in the Programme for International Student Assessment (PISA) 2018, Brazilian advantaged students outperformed disadvantaged students in reading by 97 score points. In Brazil, about 1 in 10 high-achieving disadvantaged students — but 1 in 25 high-achieving advantaged students — does not expect to complete tertiary education [9]. The drivers of the low educational level in Brazil reflect the countries’ historical neglect for investments in this area, but also to the people’s discouragement to learning *per se*. Both the neglect and people’s discouragement to learning might be related to the increasing deindustrialization and intensification of a neoextractivist economy, implemented in the peripheral capitalist countries [10, 11]. In a nutshell, while access to university had increased in Brazil, education quality is still a burning issue [10]. To promote an effective change in education, we need not only content-oriented teaching, but most importantly, a teaching for higher levels of thinking [12], allied to the broadening of access to higher education.

### Effective learning in large classes

A collateral effect of broadening access to higher education in Brazil was the increase in the number of students per class [13]. Teachers face major challenges regarding how to improve learning efficiency in ever-growing class sizes, with this issue embodying a vast range of courses and student interests [14]. The challenge is particularly acute in the context of Brazilian university education, due to increases in the diversity of the socioeconomic and cultural backgrounds of the students [5] following recent (2007 — 2014) policies concerning affirmative action and broadening access to university for low-income students [15]. Such policies have increased the proportion of students arriving from public high schools, which generally tend to have lower teaching standards [16].

Although teachers’ teaching skills may be affected by a large range of aspects, such as their lives in context [17], discourse delivery tasks [18] or pedagogical interventions [19], active learning has been shown to enhance learning effectivity in a large spectrum of university courses [20–22]. Active learning has been particularly efficient for minority students [23]. However, evidence regarding which active learning approaches result in more effective learning remains scant (but see [24], [20] and [21], and references therein), meaning we lack understanding regarding how students themselves perceive and evaluate the effectiveness of these methods.

### Transformational learning

Although all forms of learning involve a transformation of the learner’s mind [25], several learning theorists distinguish informative learning from transformational learning [26]. For this study, we favor the distinction between transformational *versus* informative learning — in Paulo Freire’s terms, problematizing or liberating *versus* sitting, or traditional, learning [27] — because we aim to analyse a form of learning designed to develop critical thinking [28]. This is particularly relevant in countries with high economic inequalities and low overall education standards, as in the case of Brazil [29]. Transformational learning involves several alternative pedagogies, such as active learning, student-centered learning, collaborative learning, experiential learning, and problem-based learning [30].

A complementary view on transformational teaching is Pierre Bourdieu’s [31] view of cultural capital as a tool for domination in the constant struggle among social classes. In expositive, lecture-based classes, the teacher, the holder of the dominant cultural capital, represents a source of knowledge, and the mainly oral transmission may reinforce cultural barriers, such as sophisticated vocabulary, which obstruct the learning of students. We propose that Paulo Freire’s problematizing education [27] is, unwittingly, a way to counter the domination promoted by lecture-based classes.

Since Bruner [32] concluded that knowledge discovered by children themselves is more prone to be used and retained than facts that are designed to be memorized, it has been suggested that active learning tasks would offer a more efficient approach to fostering learning. Despite the accumulating levels of data on the efficiency of active learning methodologies, most university and high school classes are predominantly lecture-based, focusing mainly on the teachers’ speech. To overcome the role of teachers as the purveyors of knowledge, the teacher has to problematize and provoke curiosity, doubt, and critical thinking, rather than merely being the “source of knowledge” [33]. Through such an approach, the teacher is a mediator, helping to construct and reinforce disputing points of view [34].

### Scientific teaching

Handelsman et al. [35] published a plea for “scientific teaching”, in which teaching is approached with the same rigor as science at its best, involving active learning tasks and teaching methods that have been systematically tested and shown to reach diverse student. Our work contributes to these debate, by testing active learning tasks’ efficiencies. A landmark among these studies is the meta-analysis of Freeman et al. [20], showing that active learning increases students’ performance in science, engineering, and mathematics. Examples are Fu et al.’s [36] work on writing performance, Zhang et al.’s [19] on promoting intercultural competence, and Huang’s [37] on new pedagogical methodologies to enhance critical thinking skills and creativity. The use of more sophisticated statistical analyses is proving efficient [38], distinguishing the impact of an instructional intervention from the impact of student characteristics, thus helping to elucidate the mechanisms and effectiveness of active learning.

### Aims

Most studies on active learning focus on students’ performance [20, 36, 39], or on students’ perspective [21], but we are not aware of studies comparing these two perspectives. Evaluation of the efficiency of active learning by the students themselves faces widespread suspicion concerning the accuracy of students’ self-evaluation still haunts the academic community [40], meaning there is a clear demand for data that evaluates the correlation between self-evaluation and actual exam grades. Here we contrasted both the students’ perspective and the perspective of students’ exam grades, and found that these perspectives converge.

In this study, we aimed to evaluate the effectiveness of active learning for ecology teaching, the extent to which students’ characteristics altered these effects, and the degree to which students detected the effectiveness of active learning. Our main hypothesis was that active learning increases the efficiency of ecology teaching. A complementary hypothesis was that students perceived which active learning tasks where more effective.

## Materials and methods

### License of Human Ethics Committee

The present study is part of the project registered on the Brazil Platform website, under the title “In search of effective learning: evaluating the efficiency of alternative strategies in the teaching of Ecology”, with the number CAAE 50091415.9.0000.5153. We followed all procedures required by law, including approval by the Research Ethics Committee (CEP) and the signing of the TCLE (Free and Informed Consent Term) by all students involved in data collection.

### The course of Basic Ecology

The datasets featured in this study (datasets available at S1 Dataset and S2 Dataset) refer to a one-semester course on Basic Ecology (BIO 131) offered to 176 students across 19 undergraduate programs at the Federal University of Viçosa, Viçosa, Minas Gerais state, Brazil. The course took place during the second school semester of 2015. The undergraduate programs encompassed several subject areas, ranging from biology and agronomy to civil and electrical engineering. The course was offered to three teams of students, each consisting of around 60 students. Each team attended one single lesson (50 min) and one double lesson (100 min) per week, totaling 150 minutes of lesson-time (classes) per week for each team, over 18 weeks. The following active learning tasks were undertaken, for all students. All students in this study experienced a similar amount of active learning. Therefore, what we tested was how the students’ perception of the effectiveness of the active learning tasks compare to the correlations of the students’ performance and effort in the active learning tasks with their performance in two exams.

### Evaluation of learning

To evaluate the amount of learning by the students, we used two perspectives:

- Students’ perspectives of the effectiveness of the active learning tasks in relation to their self-evaluated learning;
- the correlation of students’ effort and performance in the active learning tasks with their grades in exams.

The first approach allowed us to observe how the students themselves sensed their learning. The second approach allowed us to observe students’ ability to answer objective questions on the course’s contents, including mastering of ecological theory and scientific methodology, application of the theory to practical situations, and interpretation of graphs.

### Perspective of the students

To estimate the perspective of the students, we supplied an anonymous survey, using Google^®^ forms (complete survey available at S1 Text), which was made available for students one month before the end of the academic semester. The students could choose a grade from zero to five for each item of the self-evaluation survey. We used this survey to estimate self-evaluated learning, self-evaluated effect of active learning tasks and teachers’ stimuli for students’ active involvement, and student’s characteristics (see Explanatory variables). Raw data of students’ answers to the anonymous survey results is provided in the supporting information (S1 Dataset).

### Exams

During the 10^th^ and 17^th^ teaching weeks, the students undertook two exams. The first exam (available at S2 Text) consisted of 15 questions, of which 13 had five true or false statements each, and two questions were graphical tasks. The second exam (available at S3 Text) consisted of 12 questions, of which 11 had five true or false statements each, and one questions contained three true or false statements and two graphical tasks. Raw data on the exams’ grades is provided in the supporting information (S2 Dataset). We used exams to understand student learning more directly, independently of the students’ point of view.

### Applied active learning activities

Most active learning activities were modified after Richard Felder’s recommendations [41, 42]. We promoted five distinct active learning activities:

- Teachers’ attempts to stimulate students’ active involvement;
- Extra-class group task;
- Online writing assessments;
- Redoing exam in pairs;
- Classroom writing group assessments.

These five active learning activities were implemented for all students. We used students’ responses to these activities, to better understand how they impact the students’ experience in our ecology courses. Further, we compared the effects perceived by the students with the effects detected in correlations of students’ effort and performance in the active learning tasks with their grades in two exams, so as to evaluate in how far students’ perception coincides or differs from their actual learning. Our assumption was that students’ performance in the exams reflects their actual learning, although we are aware that this assumption is not indisputable.

### Teachers’ attempts to stimulate students’ active involvement

Along the basic ecology course, that encompassed one double (100 min) and one single lesson (50 min) per week (see The course of Basic Ecology for more details), the weekly double lessons consisted of traditional lectures, using data-show projection, supported by one to four quick active-learning activities (maximum 5 min). For the purposes of this study, active-learning activities involve tasks that: (i) stimulate students to ask questions, emphasizing that there are no “stupid” questions nor “stupid” answers, (ii) stop the lecture at least twice each class in order to ask questions directed at a single student, chosen arbitrarily from different spatial regions of the classroom, prizing both answers and doubts, and stimulating colleagues to help and complement the answer; (iii) tackle questions in groups of students through discussions of at least three-minutes within the groups. Teachers’ stimuli for active involvement do not constitute pure active learning tasks, and were not subject of students’ performance nor effort estimation, but our hypothesis is that teachers’ behaviour could contribute to students’ active involvement and, hence, learning efficiency. These attempts contrast to the predominance of traditional “sage on a stage” mode of lecturing in Brazilian universities, in all fields of knowledge, including both STEM (Science, Technology, Engineering and Mathematics) and human sciences (Arts, Philosophy, Architecture, among others) [43]. Specifically within environmental courses in Brazilian universities, active learning is restricted most often to separate presentations done by the students (called “seminars”), scattered within predominantly traditional lectures presented by the teachers [44], thus, comparatively much less active learning than implemented here.

### Extra-class group task

In this active learning task, that was carried out along the whole academic semester, the students had to work in groups of up to five members, organized by themselves, to produce a presentation to their classmates at the end of the semester. This work was undertaken outside the classroom, with an open theme and format that could consist of whatever students proposed. The accepted formats were banner, data-show presentation, research on a scientific question, lecture to high school or elementary school students, public opinion poll, video, poetry, theater presentation, song, scale model, and pedagogical installation. Supervision of the extra-class group task was done through regular tasks designed to be submitted online. These included submissions on: the theme of the extra-class group task (4^th^ week), (ii) the goals the students expected to achieve, the ecological theory involved, and the schedule and division of tasks among group members (6^th^ week), (iii) a report on progress and the contribution of each group member to the project, along with restructuring and changes to the original project that were undertaken by the students (8^th^ week). The groups scheduled their presentations using a common spreadsheet, with each presentation evaluated by the teacher and at least two student volunteers. On the day of the presentation, each student delivered a sealed envelope with grades for each member of their group, including self-evaluation (= reciprocal evaluation). Thus, we expected that members that were considered idle would receive a lower grade than the other members of the group. This strategy was designed to penalize profiteers and prize pro-active and cooperative students. The final grade for the extra-class group task was the average of the reciprocal evaluation grades, given by the colleagues within each group, multiplied by the average of the grades given by the teacher and volunteer students for the presentation. Grades for the extra-class group task could amount to up to 15 points.

### Online writing assessments

We implemented four online writing assessments, in which students had to answer two to three questions with up to 200 words. The deadline for the assignments was two weeks. To estimate students’ performance in this active learning task, each exercise was graded from 0 to 1.5, summing up to 6 points. To avoid redundancy between the estimate regarding performance and that regarding effort, we used the mean grade of the answered exercises as an estimate of each student’s performance in the online writing assessments, while student’s effort in this learning task was estimated by counting the number of answered exercises, varying from 0 to 4.

### Redoing exam in pairs

To foster cooperative learning, we implemented the task of redoing exam in pairs [45]. The first exam was resubmitted to the students two weeks after they first undertook it, with the students asked to answer the questions again in pairs. Pairs that answered more than 80% of the same questions correctly received a bonus of five points on their final grade.

### Classroom writing group assessments

Along the basic ecology course, that encompassed one double (100 min) and one single lesson (50 min) per week (see The course of Basic Ecology for more details), in the weekly single lesson, the students received a task that was designed to be discussed and answered during the class (50 to 60 min) in groups of four to five students, with three to five questions on the subject having been discussed in the previous double lesson. The total number of classroom writing assessments was six, but the final grade consisted only of the five best grades. The answers were corrected by undergraduate tutors. To estimate students’ performance in this task, each exercise was graded from 0 to 3, summing up to 15 points. To avoid redundancy between the estimate regarding performance and that regarding effort, we used the mean grade of the accomplished classroom assessments as an estimate of each student’s performance in this task, while student’s effort in this task was estimated by counting the number of accomplished tasks, varying from 0 to 6.

### Hypotheses and their predictions

We tested the hypothesis that active learning increase students’ learning, comparing two approaches: from the perspective of the students, established through self-evaluation (*n* = 86, see Fig 1 for the flowchart of the hypotheses from the perspective of the students), and from the perspective of the students’ grades (*n* = 176, see Fig 2 for the flowchart of the hypotheses from the perspective of the students’ grades in the exams). If our hypothesis was true, we expected that both approaches would agree. If students perceived that active learning tasks increased their learning, their self-evaluated learning should increase with their self-evaluated effect of active learning tasks. If active learning tasks increased students’ learning, students’ actual grades should, accordingly, increase with students’ performance and effort in the active learning tasks. If the result agreed, this would show that the students’ perception of the effectiveness of active learning tasks and of their own learning mechanisms corresponds to actual learning, as estimated by external evaluation tools.

**Fig 1.**
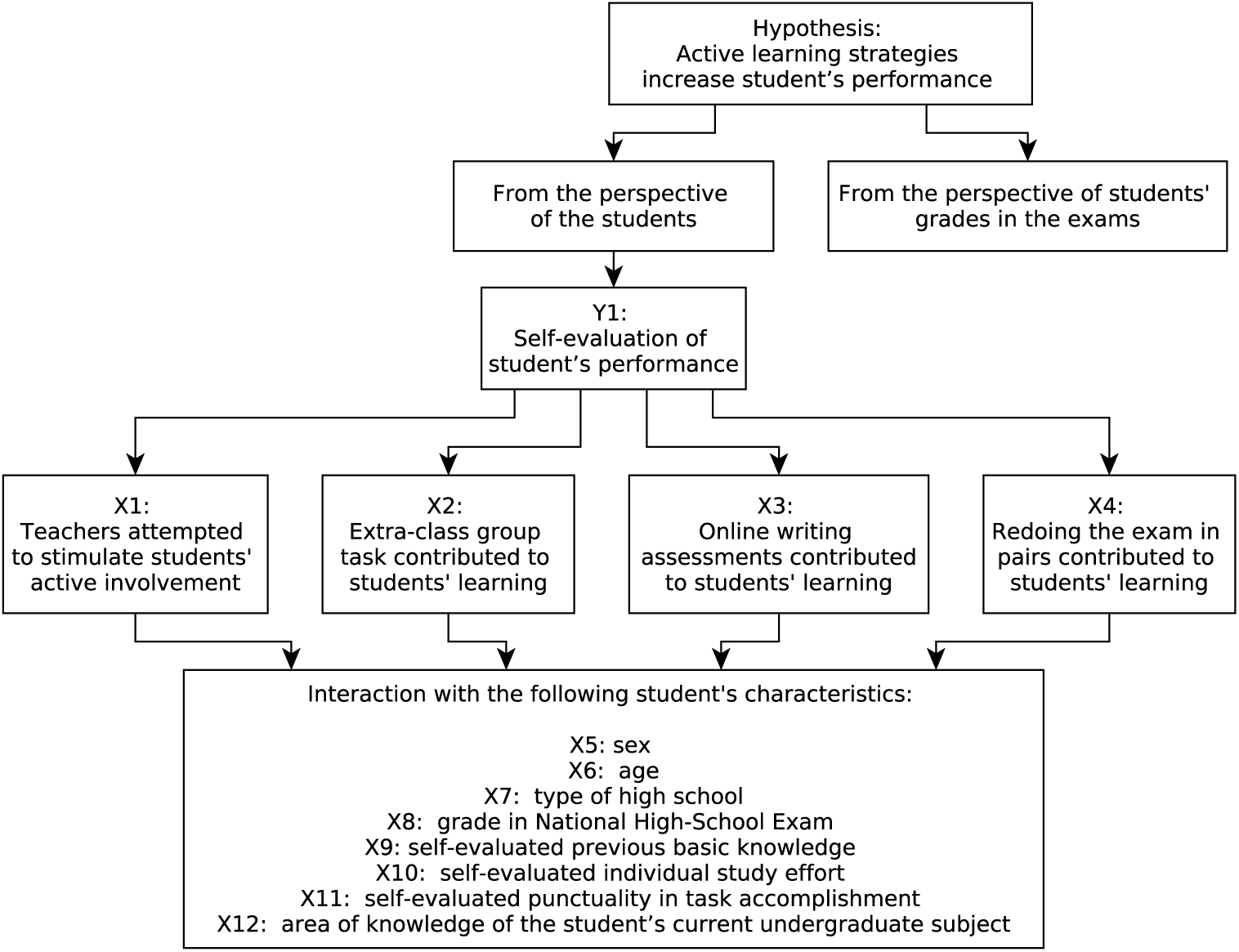
Flowchart of the tested hypotheses from the perspective of the students: active learning increases students’ performance in the basic ecology course, from the perspective of the students. See Fig 2 for the flowchart of the tested hypotheses from the perspective of the students’ grades in the exams.

**Fig 2.**
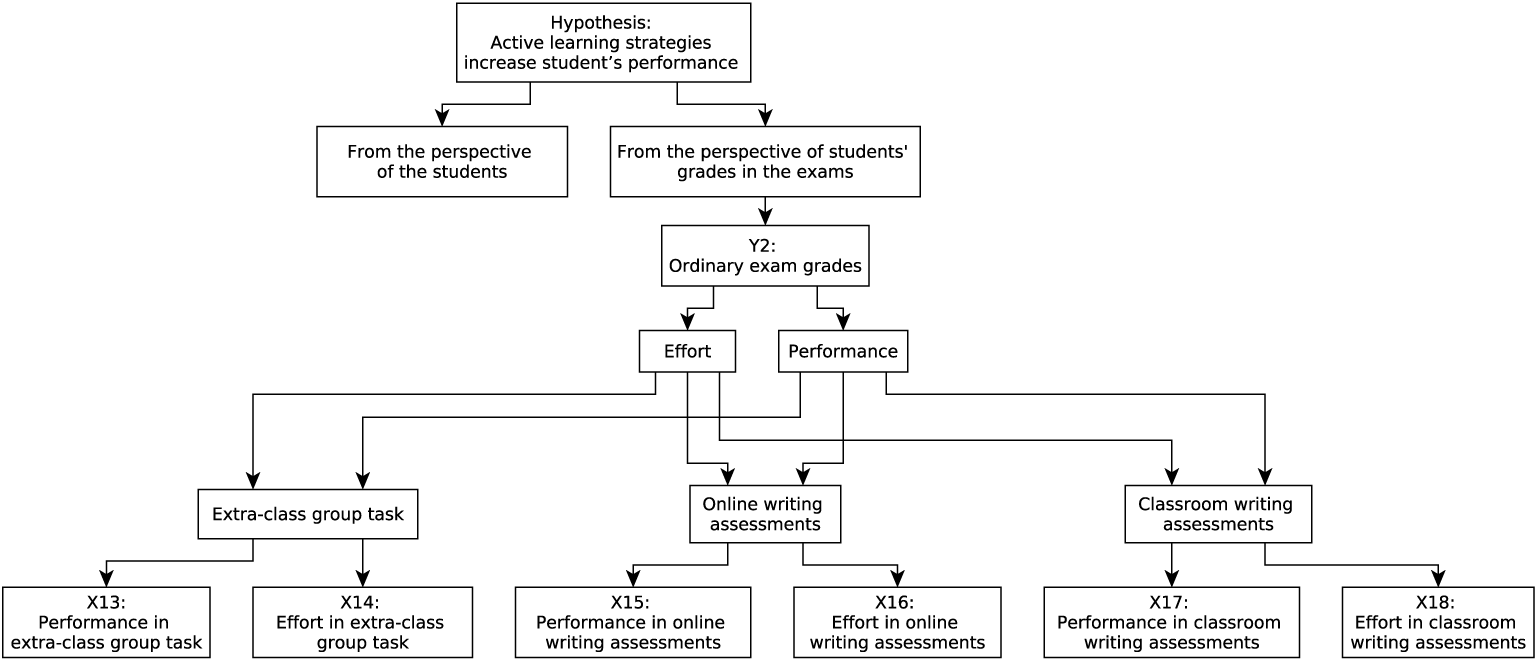
Flowchart of the tested hypotheses from the perspective of the students’ grades in the exams: active learning increases students’ performance in the basic ecology course, from the perspective of the students’ grades in the exams. See Fig 1 for the flowchart of the tested hypotheses from the perspective of the students.

If only self-evaluated performance increased with students’ perception of the effectiveness of active learning tasks, this would indicate that students’ perception might rather related to motivation or enthusiasm than effective learning. Alternatively, external evaluation tools, particularly exam grades, could be misleading, unable to detect students’ learning.

If, on the other hand, only exam grades increased with active learning tasks, this would reveal that students are unable to evaluate their own learning. Eventual correlations in the self-evaluation questionnaire would be related to self-cognition, unrelated to efficiency in answering formal exams. Such result would shed suspicion unto examination tools themselves, or unto students’ self-consciousness. Our assumption was that both students’ perceptions and their exam grades, would reflect the actual learning results.

For the perspective of the students, we were able to evaluate if the following student characteristics affected the effectiveness of active learning tasks:

- Sex;
- Age;
- High school;
- National High School Exam;
- Previous basic knowledge;
- Regular previous study;
- Punctuality;
- Area of knowledge of their undergraduate program.

### Our variables

The present work has a quantitative approach, reflecting our research expertise and experience in ecology, within the hypothetical-deductive science paradigm [46]. To carry out studies within the hypothetical-deductive paradigm it is necessary to quantify and test explicit hypotheses, using statistical analyses, so as to evaluate the null hypothesis that the working hypotheses are wrong, and all observed variation may be due to chance alone. For this, one has to translate observations into numbers, i.e., quantify them. Each of the quantified aspects is called a “variable”. The first step to evaluate hypotheses statistically is to understand which of the variables are response variables and which are explanatory variables [47].

### Response variables

The response variable is the thing we are working on: it is the variable whose variation we are attempting to understand. This is the variable that goes on the Y axis of the graph (the ordinate) [47]. In this study, we had two response variables, each one used to test one set of hypotheses:

- Self-evaluated students’ learning; and
- Students’ exam grades.

### Self-evaluated students’ learning

In order to estimate the self-evaluated students’ learning (Y_1_), we multiplied the grades that students awarded themselves concerning “motivation” (*m*), by “satisfaction with own performance” (*s*), plus the grades concerning the degree to which “the aims of the course were reached” (*a*); Y_1_ varied from zero to 30 (Eq 1). 

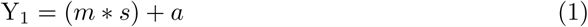

Our reasoning for multiplying the first two terms was that, for an unmotivated student (i.e., with low grades for “motivation”), low grades might have been evaluated as sufficient or satisfactory (i.e., high grade for “satisfaction with own performance”), while for more motivated individuals, satisfaction would imply higher grades. Therefore, low grades for “motivation” reduced the importance of higher grades for “satisfaction”, while high grades for “motivation” increased the importance of “satisfaction” grades. We considered the third term, related to the course’s success, as a factor independent of the student’s motivation and satisfaction levels. Our assumption for adding the grade for each student’s evaluation of the course’s success was that, independent of a student’s motivation or satisfaction, students would evaluate the course’s success equitably. We aggregated these three terms in a single response variable (Y_1_ — self-evaluated students’ learning) in order to reduce the risk of a type I error, which is to reject a true null hypothesis.

We used “self-evaluated students’ learning” (Y_1_) as the response variable to test our hypothesis from the perspective of the students (Fig 1). The summed punctuation of students’ actual grades in the two exams was the response variable (Y_2_) that we used to test our hypothesis through the students’ actual grades in the exams (Fig 2), varying from zero to 60.

### Students’ grades in the exams

Students’ exam grades (Y_2_) were a sum of their grade in each of the two exams (see Exams for details on the applied exams), each exam counting up to 30 points, so that students’ exam grades could vary from 0 to 60 points.

### Explanatory variables

The explanatory variables refer to the explanations for the response variable: they refer to the mechanisms that drive the thing you want to explain, i.e., they would affect your response variable. These are the variables that go on the X axis of the graph (the abscissa) — each explanatory variable might be plotted in a different graph, maintaining the same response variable in the Y axis. We are interested in the extent to which variation in the response variable (Y) is associated with variation in the explanatory variable (X) [47].

We had two sets of explanatory variables:

- one set of variables from the perspective of the students, and
- one set of variables from the perspective of the students’ grades in the exams.

We used 12 variables (X_1_ to X_12_) related to the hypotheses from the perspective of the students (see Fig 1 for the flowchart of the hypotheses from the perspective of the students). We used six variables (X_13_ to X_18_) related to the hypotheses from the perspective of the students’ grades in the exams (see Fig 2 for the flowchart of the hypotheses from the perspective of the students’ grades in the exams).

### Explanatory variables from the perspective of the students

The first four explanatory variables, from the perspective of the students, estimated students’ evaluation of the effectiveness of the following active learning activities:

- X_1_: “Students evaluated that teachers stimulated active involvement”;
- X_2_: “Do you consider that doing the extra-class group task contributed to your learning?”;
- X_3_: “Do you consider that online writing assessments contributed to your learning?”;
- X_4_: “Do you consider that redoing exam in pairs contributed to your learning?”.

X_1_: “Students evaluated that teachers stimulated active involvement”, was estimated by adding students’ answers to the questions “Have teachers encouraged questions?”, “Have teachers sought active involvement of students beyond simple questions?”, “Have teachers payed careful attention to student comments, questions and answers and respond constructively?”, and “Have teachers checked periodically if the students are understanding?”. X_2_ was the students’ answer to the question “Do you consider that doing the extra-class group task contributed to your learning?”. X_3_ was the students’ answer to the question “Do you consider that online writing assessments contributed to your learning?”. X_4_ was the students’ answer to the question “Do you consider that redoing exam in pairs contributed to your learning?”. Students had to rate these questions with integer values, meaning that the explanatory variables ranged as follows: X_1_ from 0 to 20; X_2_, X_3_ and X_4_ from 0 to 5.

We used further eight explanatory variables to evaluate students’ individual characteristics:

- X_5_: “sex” (male, female, other);
- X_6_: “age” (*<* 19, 19 to 24, *>*24);
- X_7_: “type of high school training” (private school, state public school, federal public school, Agricultural Family School (EFA – *Escola Família Agrícola*));
- X_8_: “grade in the National High-School Exam” (Exame Nacional do Ensino Médio — ENEM; 1 to 4 ^2^;
- X_9_: “self-evaluated previous basic knowledge” (“Did you have the basic training necessary to achieve good results in the course?”, 1 – 5);
- X_10_: “self-evaluated individual study effort” (“Have you studied regularly and in advance the content presented?”, 0 – 4);
- X_11_: “self-evaluated punctuality in task accomplishment” (“Have you done the requested activities on time? “, 0 – 5); and
- X_12_: “area of knowledge of the student’s current undergraduate program” (agrarian, biological, exact, human).

These eight variables (X_5_ to X_12_) were extracted from the self-evaluation survey (complete survey available at S1 Text). We evaluated if any of these characteristics of the students affected effectiveness of active learning tasks (see Fig 1 for the flowchart of the hypotheses from the perspective of the students).

### Explanatory variables from the perspective of the students’ grades in the exams

In order to test our hypothesis from the perspective of the students’ grades in the exams (see Fig 2 for the flowchart of the hypotheses from the perspective of the students’ grades in the exams), we used six variables (X_13_ to X_18_, related to three active learning activities:

- Extra-class group task;
- Online writing assessments; and
- Classroom writing assessments.

We were able to distinguish between the impact of students’ **performance** regarding the task (represented by their grade in the task) and students’ **effort** in that task (represented by the number of intermediate steps completed by the student for each task). Thus, for each of these activities, we used two aspects:

- students’ performance in the task, and
- students effort in the task.

Effort relates to how much each student invested in coping with the task. Some students might have skipped one or two intermediate steps, missing to answer them in the online platform, but even so, completed the task, therefore being evaluated by the teacher in their performance in the task. Other students might have done all steps to complete the task (which we interpreted as maximum effort), whilst the end result (the grade attained in the task), i.e. performance in the task, was not optimal. Performance was the result of the students’ investment, translated into the grades the student attained in the task. Performance in each task was estimated by the respective grade. Effort in each task was estimated by the proportion of completed steps involved in the task.

To guarantee independence between the estimates of a student’s performance and effort in each task, estimates of the performance of each student excluded steps that were not answered by this student in the task, calculating the average grade of the answered tasks. For example, in the task “online writing assessments”, there were four assessments along the teaching semester. If a student answered to only three of these, his/her effort was equal to 3/4, while his grade in the task was the average of his/her grades in the three answered assessments. Thus, we guaranteed that even low-effort students, that answered a low proportion of the tasks, could have high performance estimates.

The explanatory variables for students’ exam grades (see Fig 2 for the flowchart of the hypotheses from the perspective of the students’ grades in the exams) were:

- X_13_: “performance in the extra-class group task”;
- X_14_: “effort in the extra-class group task”;
- X_15_: “performance in the online writing assessments”;
- X_16_: “effort in the online writing assessments”;
- X_17_: “performance in the classroom writing assessments”; and
- X_18_: “effort in the classroom writing assessments’.

X_13_: “performance in the extra-class group task” corresponded to the students’ grades in this task, varying from zero to 15. X_14_: “effort in the extra-class group task” corresponded to the number of the regular tasks submitted online by each student, varying from zero to 6. X_15_: “performance in the online writing assessments” corresponded to the average grade of the answered assessments, varying from zero to 6. X_16_: “effort in the online writing assessments” corresponded to the number of answered assessments, varying from 0 to 4. X_17_: “performance in the classroom writing assessments” corresponded to the average grade in the answered assessments, varying from zero to 15. X_18_: “effort in the classroom writing assessments’ corresponded to the number of accomplished assessments, varying from zero to 6.

## Statistical analyses

To test our hypothesis from the perspective of the students, we used analyses of co-variance (ANCOVA), with self-evaluated students’ learning (Y_1_) as response variable, all quantitative and categorical explanatory variables (X_1_ to X_12_) and the two-level interaction terms of students’ characteristics (X_5_ to X_12_) with their evaluation of the effectiveness of the active learning tasks (X_1_ to X_4_), in generalized linear models (GLMs) with normal distribution. Therefore, the complete model for self-evaluated students’ learning included all four terms of students’ evaluation regarding the effects of the active learning tasks (X_1_:”teachers stimulated active involvement”, X_2_:”the extra-class task”, X_3_:”redoing exam”, and X_4_:”online writing assessments”), the eight students specific characteristics (X_5_ to X_12_), and the interaction terms of each characteristic with each active learning task: 

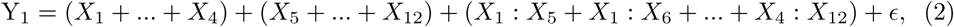

where “*X*_1_: *X*_5_” stays for the two-level interaction of the terms *X*_1_ with *X*_5_, and so forth, while “*ϵ*” stays for the random error, with normal distribution. See Explanatory variables from the perspective of the students for explanation of each term.

As a result, there were 12 terms for each explanatory variable and 32 second-level interaction terms in total. The interaction terms enabled us to evaluate the extent to which separate active-learning tasks interacted differently with the specific characteristics of the students.

To test our hypothesis from the perspective of the students’ grades in the exams, we used multiple regression, with students’ exam grades as response variable (X_2_) and students’ performance and effort in the active learning tasks (X_13_ to X_18_) as explanatory variables, adjusting GLMs with normal distribution: 

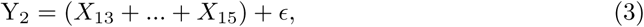

where “*ϵ*” stays for the random error, with normal distribution. Here we did not include any interaction term, because we had not the characteristics of the students for these data-set. See Explanatory variables from the perspective of the students’ grades in the exams for explanation of each term.

We evaluated nonlinearity for each complete model by adjusting generalized additive models (GAMs) with normal distribution, using “mgcv” package [48], including a smoother for each explanatory variable relative to active learning tasks. GAM is a generalized linear model in which the linear predictor depends linearly on unknown smooth functions of some predictor variables, and interest focuses on inference about these smooth functions. The smooth function is a curve adjusted to the data, that can vary from linear, in which case the estimated degrees of freedom (edf) is equal to one, to curvilinear, in which case the edf is higher than one. We used the “gam.check” procedure [49] to evaluate if the adjusted GAM’s basis dimension (*k*) were adequate. We used *F* test (procedure *anova* of the adjusted GAM model) to evaluate if the adjusted curves presented edf higher than 1 (*edf >* 1), and plotted the adjusted GAM curves to evaluated if they presented non-linear shape [50]. Explanatory variables that presented evidence of nonlinearity were maintained as non-linear smoother, i.e., the curve adjusted by the GAM, while explanatory variables with *edf* = 1 and linear shape were adjusted with linear predictor, i.e., linear regression. If nonlinearity was detected, we used the estimated non-linear effects of each explanatory variable in the minimum adequate model, summed to the linear effect of those variables where there was no non-linearity detected. Finally we compared the models with and without smoothers using ANOVA. If there were significant differences, we chose the model with lowest Akaike Information Criterion (AIC) [51].

We tested collinearity of the explanatory variables using individual multicollinearity diagnostics with the “imcdiag” function of the package “mctest” [52, 53]. Pairs of explanatory variables that had a correlation value *>* 0.7 were considered collinear. In that case, we adjusted the complete model excluding each of the collinear explanatory variables separately, and chose the model with the lowest AIC value. We also tested the effects of suspected outliers by detecting them in the plots using the adjusted curve, withdrawing them from the data, adjusting the same model, and comparing the results.

If outlier removal altered the predictions qualitatively (i.e., changing the significance or direction of the effect), we deleted it.

Significance of the explanatory variables was evaluated by deletion of non-significant terms, beginning with interaction terms [51]. Therefore, complete models were simplified until the minimum adequate model (MAM) was achieved for each response variable.

To draw the graphs of the MAM, we plotted the observed values for each of the response variables (*Y*_1_ = self-evaluated students’ performance; *Y*_2_ = ordinary exam grades in the Y axis, and the observed values of each of the significant explanatory variables (*X*_*j*_) in the X axis. To draw the adjusted curves of the MAM, we calculated the effect of each significant *X*_*j*_ on *Y*, in the scale of the observed Y values, as bellow:

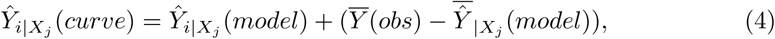

being 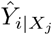 (*curve*) the value that we used to draw the adjusted curve/straight line in the graph; *X*_*j*_ each of the explanatory terms of the MAM; 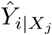 (*model*) the estimated value for the effect of the *X*_*j*_ explanatory term, taken from the MAM object; 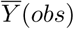 the overall mean of the observed *Y* value; 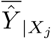 (*model*) the mean of the estimated values for the effect of the *X*_*j*_ explanatory term, taken from the MAM object.

We discarded students’ surveys where not all questions evaluated in our hypothesis were answered. We presented exact *P* values, or, when referring to more than one result, the highest *P* value when the set of results was significant (i.e., *P <* 0.05), or the lowest *P* value, when the set of results was non-significant (*P ≥* 0.05). Adjusted models were subjected to residual analyses. All analyses were done using R [54].

## Results

Here we present a brief recap, showing the hypotheses that we tested and summarizing the main results. Afterwards we present the results formally, with their respective statistics. To evaluate our main hypothesis that active learning increases the efficiency of ecology teaching, we compared two approaches: (i) from the perspective of the students’ and (ii) from the perspective of students’ grades in the exams (see Hypotheses and their predictions for more details). According to the first approach, our hypothesis was that students perceive that active learning tasks increase their ecology learning. If this were true, we expected that students’ self-evaluated learning would increase with students’ perception of the effectiveness of the active learning tasks. Within this approach we also evaluated if students characteristics, such as sex, age and previous knowledge, affected the effectiveness of active-learning tasks. If any of these characteristics affected the efficacy of active learning tasks, we expected that there would be a significant interaction of the characteristic with the effect of the affected active learning task upon students’ learning. An interaction between two explanatory terms means that the effect of students’ perception on their learning would differ among students with different characteristics, such as age.

We accepted the hypothesis of our first approach, that students’ self-evaluated learning increased with their evaluation of active learning efficacy, for three aspects of active learning (Fig 3A-C):

**Fig 3.**
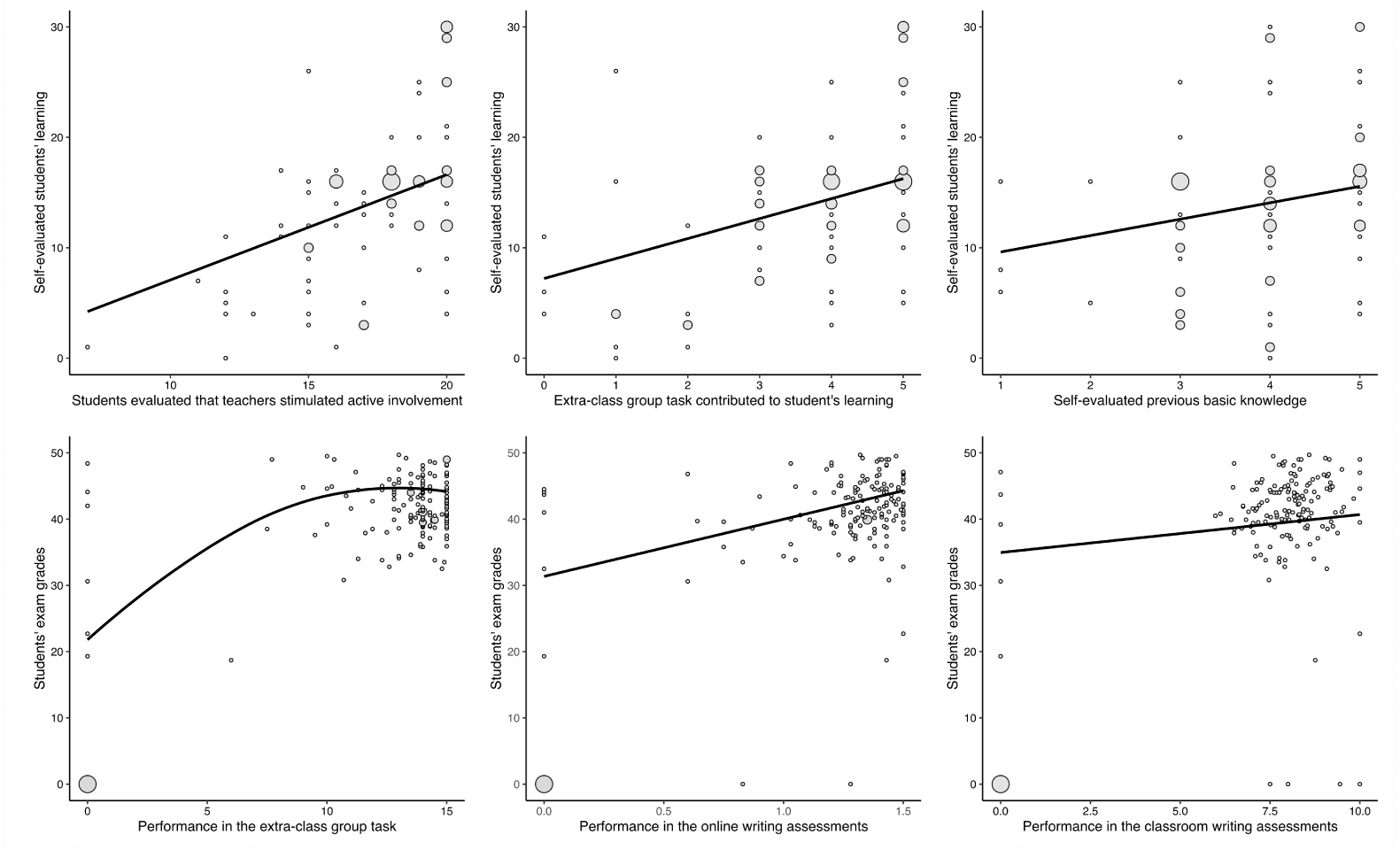
Effectiveness of active learning on students’ performance in ecology: the perspective of students *vs* actual grades. This figure shows which explanatory variables (related to active learning) were correlated with which response variables (see Our variables for more details). (**A**) Students’ self-evaluated performance increased linearly with students’ evaluation of the effectiveness of teachers’ stimuli for active involvement. (**B**) Students’ self-evaluated performance increased linearly with students’ evaluation of the effectiveness of extra-class group task. (**C**) Students’ self-evaluated performance increased linearly with students’ evaluation of their previous basic knowledge. (**D**) Students’ grades in exams increased asymptotically with their performance in extra-class group task. (**E**) Students’ grades in exams increased linearly with their performance in online writing assessments. (**E**) Students’ grades in exams increased linearly with their performance in classroom writing assessments. Larger circles indicate multiple overlapping points.

- teachers’ stimuli for students active involvement,
- extra-class group task, and
- students’ previous basic knowledge.

From the students’ perspective, the other two active learning tasks that we tested with these data, online writing assessments and redoing exam in pairs, were not correlated to students’ self-evaluated ecology learning.

According to the second approach, from the perspective of students’ grades, our hypothesis was that students’ grades increase with their performance and effort in the active learning tasks. Within this approach, we evaluated two aspects of the active learning tasks: students’ performance and students’ effort in the task.

We accepted our hypothesis, from the perspective of students’ grades, for students’ performance in all tested active learning tasks (Fig 3D-F):

- extra-class group task,
- online writing assessments, and
- classroom writing assessments.

There was no effect of students’ effort in the active learning tasks on their grades in exams, either because effort was collinear with performance in the task (extra-class group task and online writing assessments), or because the effect was not significant (classroom writing assessments). The results of both perspectives agreed, showing not only that active learning tasks were effective, but also that students’ perception of the effectiveness of active learning tasks and of their own learning mechanisms is adequately estimated by external evaluation tools.

### Active learning tasks increased students learning, from the perspective of the students’

We did not detect any collinearity among the explanatory variables for self-evaluated students’ learning (all r values *<* 0.65), meaning that none of the explanatory variables was redundant. Students’ self-evaluated performance increased linearly with their evaluation of teachers’ stimuli for students’ active involvement (*F*_1,77_ = 14.031, *P* = 0.0003454, Fig 3A), with extra-class group task (*F*_1,77_ = 13.987, *P* = 0.0003522, Fig 3B) and with previous basic knowledge (*F*_1,77_ = 5.6441, *P* = 0.02, Fig 3C). There were no significant interaction terms (*F <* 1.3, *P >* 0.3), and neither online writing assessments, redoing exam in pairs, nor students’ sex, age, type of high school, grade in the National High-School Exam, individual study effort, punctuality nor area of knowledge, affected students’ self-evaluated performance (*F <* 1.3, *P >* 0.29). There was no evidence of nonlinearity (*edf <* 1.25), and the models with and without smoothers were similar (*F* = 0.921, *P* = 0.2912), with lower AIC for the model with no smoothers (*AIC* = 512.9135) than with smoothers (*AIC* = 513.1675), meaning that all tested active learning aspects affected students’ self-evaluated learning linearly.

### Active learning tasks increased students learning, from the perspective of students’ grades in the exams

We detected collinearity of performance and effort in extra-class group task (*r* = 0.704) and between performance and effort in online writing assessments (*r* = 0.707), meaning that for these two explanatory variables, the effects of students’ effort and performance upon students’ grades were redundant. Using AIC, we chose the best model for each of the collinear explanatory pair of variables. The best model included performance in the online writing assessments and performance in the extra-class group task (*AIC* = 1185.275), compared to performance in extra-class group task and effort in online writing assessments (*AIC* = 1219.823), to effort in extra-class group task and performance in online writing assessments (*AIC* = 1224.531), and to effort in extra-class group task and effort in online writing assessments (*AIC* = 1261.478). Thus, for these two active learning tasks, the effect of students’ effort was redundant with students’ performance, and performance was the best predictor of students’ grades in the exams.

There was evidence of nonlinearity for the response of students’ exam grades to students’ performance in extra-class group task (*edf* = 2.325, *F* = 31.298, *P* = 9.69^−15^), but not for the remaining explanatory variables in the full model (*edf <* 1.003, *F* = 0.0431, *P* = 0.2678). Thus, we adjusted the models using the gam procedure, used to include non-linear terms in the model, but with only one smoother for extra-class group task performance, meaning only this term was non-linear, while the remaining explanatory variables were adjusted with a linear response. The model with one smoother term, for students’ performance in extra-class group task, and two linear terms, for students’ performance in online writing assessments and for students’ performance in classroom writing assessments, presented lower AIC (1170.638) than the model with the same explanatory terms but without smoothers (*AIC* = 1183.598), meaning that the best model included a single non-linear term.

Students’ exam grades increased asymptotically with students’ performance in the extra-class group task (*edf* = 2.318, *F* = 40.25, *P <* 2^−16^, Fig 3D), and increased linearly with performance in online writing assessments (*df* = 1, *F* = 32.343, *P* = 5.5*e* − 08, Fig 3E), and with performance in classroom writing assessments (*df* = 1, *F* = 4.738, *P* = 0.0309, Fig 3F). Students’ effort in classroom writing assessments did not affect students’ exam grades (*F* = 0.7196, *P* = 0.3983).

### Agreements of the perspective of the students with the perspective of students grades in the exams

From both perspectives, active learning activities increased students’ learning (see Fig 3). Among the two active learning tasks that were evaluated in both perspectives (Extra-class work in groups and Online writing assessments), the extra-class work in groups increased both self-evaluated learning and students’ grades in the exams (see Fig 3 B, D), while online writing assessments increased students’ grade in the exams (see Fig 3 E), but its effectiveness was not correlated to students’ self-evaluated learning. The maintenance of the three explanatory variables (teachers’ stimuli for students’ active involvement, extra-class group task and previous basic knowledge) in the model for students’ perspective, and of the three active learning tasks (extra-class group task, online writing assessments and classroom writing assessments) in the model for students’ grades in exams, show that these variables were complementary to promote students’ learning.

## Discussion

Here we present a brief recap of the hypotheses that we tested, summarize the main results, and discuss why these results are important, what they imply, and what effective practices in university and high-school teaching might be introduced or reinforced based on our findings. We tested two hypotheses, to elucidate the effects of active learning tasks on students’ ecology learning. The first hypothesis was that students’ perceive that active learning tasks increase their ecology learning. Within this hypothesis we also evaluated if students characteristics, such as sex, age and previous knowledge, affected the effectiveness of active-learning tasks. The second hypothesis was that active learning tasks increase students’ ecology learning, from the perspective of students grades in exams. Within this hypothesis, we evaluated two aspects of the active learning tasks: students’ performance and students’ effort in the task.

### Students’ perception of the effectiveness of active learning

Our results evidenced that students perceive the effectiveness of active learning for (i) teachers’ stimuli for students active involvement, (ii) extra-class group task (both with a p value lower than 0.0004), and (iii) students’ previous basic knowledge (with a p value lower than 0.03). These three effects were linear, and there was no evidence of collinearity among these variables. We think this means that (i) teachers’ behavior in class, stimulating the active involvement of the students, and (ii) active learning activities that are performed in group and that stimulate independence, are of utmost importance in fostering students’ learning. The predominance of these two pedagogical aspects is reflected in the extremely low p values, showing their strength in driving students’ self-evaluated learning. We discuss these first two effects bellow. We discuss the third effect, that of students’ previous knowledge, and the absence of collinearity, after that.

We were not able to fully validate the survey and questions answered by the students, as, for example, the pioneering work on validating the students’ perspective on their engagement in active-learning of Ben Wiggins and colleagues [21]. Such validation would require a follow-up of several semesters, and extrapolates the available education atmosphere within which we work.

### Effectiveness of teachers’ behavior

The high effectiveness of teachers’ stimuli, from the students’ perspective, highlights the importance given by the students to teachers’ behavior. One of the key findings from decades of educational effectiveness research is the importance of the ‘classroom level’ as a predictor of pupil outcomes, and a large proportion of this classroom-level variance can be explained by what teachers do in the classroom [55]. Teachers’ attempts to elicit students’ engagement is knowingly effective [18]. In our study, teachers’ actions were to (i) stimulate students to ask questions, (ii) emphasize that there are no “stupid” questions nor “stupid” answers, (iii) challenge the students during the class, by asking questions directly to single students, (iv) prize both answers and doubts, (v) stimulate colleague students to help and complement the answer; and finally (vi) introduce active learning dynamics through questions that should be answered in little groups of students through discussions of at least three-minutes (such as recommended by Felder & Brent [41] — see Teachers’ attempts to stimulate students’ active involvement for more details). Our main goals with this strategy were to elicit critical thinking and respect and appreciate students individually. Critical thinking was already shown to be an efficient tool for teaching effectiveness [56].

Teachers’ stimuli for the students’ active involvement could be classified as a constructive activity, within the ICAP framework (**I**nteractive *>* **C**onstructive *>* **A**ctive *>* **P**assive [57], in as far as the teachers’ stimuli could boost students to synthesize their own ideas and generate a novel output, but students’ rarely responded up to this level in the classroom. Most students’ answers to teachers’ stimuli were rather on the active level, where students engage in a substantive exchange of ideas leading to a new level of understanding. Even so, our results showed that such active learning aspect is effective in boosting students’ learning.

Students’ evaluation of the effect of teachers’ classroom stimuli showed an increase in dispersion with self-evaluated performance (see Fig 3 A), meaning that there were students with high as well as low self-evaluated learning among those that evaluated teachers’ stimuli as effective, and that this variance was reduced for students that evaluated teachers’ stimuli as ineffective (note the fan-shaped dispersion of the points around the adjusted line in Fig 3 A). We interpret this result as indicating that students that had perceived themselves as having low performance (self-evaluated performance was estimated by “motivation”, “satisfaction with own performance” and “the aims of the course were reached” (see Self-evaluated students’ performance for more details), tend to evaluate that teachers’ stimuli for their active involvement were not effective — or, reciprocally, students that were not touched by the teachers’ efforts presented low self-evaluated performance. However, simultaneously, there were students with both high and low self-evaluated performance levels that rated teachers’ stimuli for their active involvement as effective. We interpret that this highlights that, contrary to the literature that states that teachers that are evaluated by the students as effective tend to be those with whom the students achieve higher grades [58, 59]. Our results indicate that teachers that require higher effort and tougher tasks, are also evaluated as more effective [60, 61], irrespective of the students’ end results. At the same time, the increase in dispersion of self-evaluated performance with teachers’ stimuli highlights the diversity of learning mechanisms: for some students, teachers’ stimuli were not effective in fostering their learning. Thus, although the average students’ learning was fostered by teachers’ stimuli for active involvement, for part of the students such stimuli do not translate into learning. This strengthens the importance of applying a diverse array of active learning tasks simultaneously.

Teachers’ stimuli for students’ active involvement go along with Paulo Freire’s [27, 62] quest for a “problematizing” education, where the teacher should not be above the students, as unique owner of knowledge, but, on the contrary, the teacher should be rather a catalyst of students’ learning. We think such behavior eases students’ insecurities and prizes students’ individuality, and, probably more important than that, fosters students’ tolerance and acceptance of differing points of view. Reciprocal respect for one’s autonomy and dignity is an ethical imperative and not a concession that we can or cannot grant each other. For Paulo Freire [33], the teacher who disrespects the curiosity of the student, his restlessness and language, violates the ethical principles of our existence. The teachers’ behavior that we considered in this work are perfectly in accord with Freire’s recommendations. Thus, our results reinforce the theory put forward by Paulo Freire [27, 62] and the constructivist school [63, 64], that learning is a common construct that must be built on personal experiences [33, 65]. In these times of intolerance, prejudice and right radicalism [66, 67], and particularly in view of the recent increase in the amplitude of students’ background in Brazilian universities (see Educational challenges in a colonized country), where, for the first time in Brazilian history, the proportion of Afro-descendant and lower socioeconomic strata students entered university [4] (but see [68] for current threats to Brazilian higher education), such fostering of tolerance and plurality is of utmost importance.

Thus, based on our results, we recommend that teachers should open themselves to students’ differences and plurality, and stimulate students’ active involvement. Treating students fairly and with respect is not only an ethical imperative, but also increases the effectiveness of learning.

### Effectiveness of the extra-class task in groups

We interpret that the extra-class task in groups (see Extra-class group task for more details) was effective because it fosters cooperative learning among students (see [69] for the effectiveness of collaborative approaches in undergraduate teaching), besides fostering independence, initiative, appreciation of the students’ particular interests, which may not be the same as the teachers’, and foster also a connection of education inside university with the society outside university. This last point was variable among groups: several groups included surveys with the population in the streets or lectures in elementary or high-school classrooms, or prepared video-presentations interviewing people in the streets, which we interpret as a way of connecting the students with the society outside the university. Other groups walked a different path, for example interviewing professors of related scientific areas in the university, or restricting their project to an internet survey and data-show presentation. These projects were less or not connected at all with society outside university. These discrepancies are part of the idea of giving freedom of choice to the students. We interpret that giving such freedom is a manner to foster confidence and responsibility upon the students. The extra-class group task is a constructivist teaching tool [64], which might catch the interest and engagement of otherwise demotivated students.

The extra-class task in groups might be classified as interactive, the highest level of learning within the ICAP framework [57], as far as in this task, students engage in a substantive exchange of ideas, which might have lead to a new level of understanding. Despite the absence of objective evaluation of in how far the students changed their understanding, our results show that this task contributes to students’ learning.

Some, albeit few, presentations of the end results of extra-class task in groups involved alternative presentation modes, such as theatre, song, scale model and poetry. We consider that opening the possibilities for such unconventional formats, constitute an important pedagogical strategy, so as to prize a broad spectrum of cognitive diversities.

Based on our results, we recommend that teachers implement extra-class group tasks in their courses, particularly if the course is in ecology, environmental or in social sciences, as a tool to foster collaborative learning and to provide space for students’ individual interests.

### Effectiveness of students’ previous basic knowledge

The third driver of students’ self-evaluated learning was previous basic knowledge, i.e., pre-college education and education outside the school. This shows that there is an important driver of learning that extrapolates university teaching. This result relates to the classic statement that the most important single factor influencing learning is what the learner already knows [70]. We see this result as a cautionary note on the limits of in how far university teaching may level differences in basic high-school education. The effect of previous basic knowledge presented the lowest slope, among the significant effects perceived by the students. This fact, added to the significance of the first two active learning approaches, shows that active learning activities are effective, albeit not overarching.

The significant effect of previous basic knowledge contrasts with the absence of learning effects related to type of high school and of grade in the National High School Exam. This may be due to the broader coverage of the National High School Exam and of high school teaching as a whole, extrapolating the subjects relevant to ecology.^3^ Basic knowledge for ecology represents only a small proportion of the whole exam.

Besides, complementary sources of knowledge, such as TV documentaries, may also contribute to the basic knowledge useful for ecology learning. Thus, students’ perception of previous basic knowledge for their ecology learning, was more accurate than the apparently objective variables such as type of high school and students’ grade in the National High School Exam. We consider that our results show that basic knowledge for ecology learning includes much more than formal high school training.

Our results evidence that teachers need be alert about the differences among student’s background. Students have different basic knowledge, and this affects their learning. Ideally, teachers should tackle this heterogeneity directly. We think, using active learning and cooperative learning techniques is an efficient way to minimize these inequalities.

### Independence of explanatory variables for self-evaluated students’ performance

The absence of collinearity among the explanatory variables for self-evaluated students’ performance demonstrates that the self-evaluated effectiveness of all evaluated active learning tasks, including teachers’ behavior, were not redundant, and that the students distinguished among activities. We interpret this as an evidence that there are actual differences in the effects of these activities depending on the student, being, therefore, complementary. A further evidence for implementing multiple active learning tasks simultaneously was the increase in dispersion of self-evaluated performance with students’ evaluation of teachers’ stimuli for active involvement, because it showed that part of the students were not affected by these stimuli. The remaining active learning tasks did not present unequivocal variation in dispersion of self-evaluated performance with students’ evaluation of the active learning task efficiency. Based on these results, we suggest that teachers should apply a plurality of active learning tasks, to maximize students’ learning.

### The active learning tasks that were not perceived as effective by the students

Our results detected no effect of students’ perception of the effectiveness of redoing exam in pairs nor of online writing assessments, on students’ self-evaluated learning. The inefficacy of redoing exam in pairs weakens Felder & Brent’s [71] recommendation, at least when applied together with other active learning activities. Redoing exam in pairs differs from the the other group tasks in as much as it restricts the “group” of students to two person, and it restricts the time available for discussion, and thus, eventual cooperative learning. Redoing exam in pairs would be classified as having a low level of active learning, according Chi & Wiley’s [57] ICAP framework. According to personal statements of some students, redoing exam in pairs was counterproductive, as far as correct answers were questioned by the partner, leading to increased errors in the answers. On the other side, we did not evaluate eventual effects of redoing exam in pairs on students’ exam grades, therefore there could be a subtle, albeit effective, increase in students’ learning fostered by this task. The absence of perceived efficacy of online writing assessments by the students, contrasts with the significant effect of students’ performance in online writing assessments unto students’ exam grades. This suggests that students underestimated online writing assessment efficacy, which might be related to this task being more tedious, and thus less appreciated by the students. The absence of effect of most students characteristics — sex, age, type of high school training, grade in the National High-School Exam, individual study effort, punctuality and area of knowledge of the student’s current undergraduate program — must be interpreted with caution, as far as our work was not designed to evaluate these effects. Probably the most interesting and surprising result was that students’ individual study effort did not affect their self-evaluated performance, but there might be a bias in these students’ evaluations.

### Effectiveness of active-learning tasks from the perspective of students’ grades in the exams

Our results showed that active learning tasks increase students’ ecology learning, from the perspective of students grades in the exams, for students’ performance in all tested active learning tasks: extra-class group task and online writing assessments, with a p value lower than 0.00001, and classroom writing assessments, with a p value lower than While the first effect, of students’ performance in the extra-class group task, was asymptotic, the two remaining effects on students’ exam grades, were linear. We detected collinearity between performance and effort in extra-class group task and between performance and effort in online writing assessments (*r* value larger than 0.7).

We interpret that the lower p values for the effects of extra-class group task and online writing assessments indicate a larger importance of these two active learning tasks on students’ learning. Overall, our results evidenced a significant and large effect of the tested active learning tasks on students’ exam grades. Besides, our results also evidenced that the tested active learning tasks are complementary, rather then redundant. We discuss these first two effects — extra-class group task and online writing assessments — bellow. We discuss the third effect, of classroom writing assessments, as well as the collinearity among explanatory variables, after that.

### The asymptotic effect of students’ performance in extra-class group task

The effect of students’ performance in the extra-class group task on students’ exam grades was asymptotic, indicating that when students achieve a performance above average, the effect of this task on learning stabilizes. Such subtlety was not detected by the students’ self-evaluation, suggesting a lower accuracy of students’ perception, compared to the effects detected in their exam grades. We interpret that the asymptotic stabilization indicates that doing the extra-class work by itself is already sufficient to enhance students’ learning, provided that the students’ performance in the extra-class work is above average. We suggest that the evaluation of performance in this task works as a driver for students’ learning and practicising cooperation.

### The linear effects of students’ performance in writing assessments on their exam grades

The linear effects of online writing assessments and of classroom writing assessments, on the students’ exam grades, show that for these active learning tasks, teacher should stimulate students’ performance maximization. Students’ training in writing is of utmost importance, not only within the ecology course, but in their formation as professionals. The effectiveness of writing tasks on students’ learning is already established [72, 73]. Our results highlight that these active learning tasks are effective and complementary, both when done in group (the classroom writing assessments) as when done individually (the online writing assessments). These two approaches have important differences: while the classroom assessment in groups requires talking and exchanging ideas among students’, boosting cooperative learning, the online assessment is answered individually, boosting learning through writing [74] — although students might have talked and exchanged ideas on these assessments as well, as far all had the same questions to be answered at home, within a two week’s period. Both writing and talking are effective learning tools, so that talk combined with writing enhances the retention of science learning over time [75].

### Complementarity of the active learning tasks

The maintenance of all three tested active learning tasks — extra-class group task, online writing assessments and classroom writing assessments — in the minimum adequate statistical model, all of them increasing students’ exam grades, shows that the effects of these tasks are complementary. Therefore, based on our results, we recommend that teachers should apply several active learning tasks altogether, so as to maximize their effectiveness on students’ learning.

### Collinearity of performance and effort

The collinearity of effort and performance in the extra-class task in groups and in the online writing assessments, evidence that for these two active learning tasks, our data where not sufficient to allow a separation of the effects of students’ effort from the effects of students’ performance, unto students’ grades in the exams. These collinearities indicate that students’ performance in these tasks correlates to their effort: students that invested a greater effort, achieved higher performance in these tasks. This correlation could be criticised as being self-evident, but previous studies found a negative correlation of students’ effort with their performance [76], which might result from the variation in students’ learning abilities. Our study did not aim to evaluate this correlation, but the results highlight that the learning mechanisms elicited by different active learning tasks may differ, as evidenced by the contrasting absence of collinearity for classroom writing assessments. Our results show that there is a contrast between the learning mechanisms of classroom writing assessments, where the effects of students’ effort and performance, unto students’ exam grades, were not collinear, compared to extra-class work and online writing assessments.

### Non-singificant effects on students’ exam grades

The single explanatory variable that was not significant to explain students’ exam grades was their effort in classroom writing assessments. Their effort was estimated by the number of assessments to which each student contributed. Each classroom writing assessment corresponded to a 50 min lesson, were the students had contribute to a written answer by a group of up to five students (see Classroom writing group assessments for further details). This result shows that, for this active learning task, the mere presence/participation in a group that answers classroom writing assessments was not sufficient to enhance learning. To increase learning, students had to take part in groups that made good answers to the assessments. Based on our results, we recommend that teachers apply writing group assessments among their active learning tasks, but also evaluate the students’ performance in this task, and include each students’ performance in the task, in the overall evaluation of students’ final grades, so as to stimulate maximization of each student’s performance in this task.

### Comparison of students’ perspective with the results of their exam grades

Overall, there was a high agreement between the effectiveness of active learning for ecology teaching through the perspective of the students, compared to the perspective of the students’ grades. The effectiveness of the extra-class work in groups was revealed in both perspectives: through students’ self-evaluation and through students’ exam grades. We interpret that this agreement is strong evidence of the accuracy of students’ perception of their learning mechanisms.

An informative aspect of the comparison between students’ perception and students’ exam grades, in relation to the extra-class group task, was that there were more students that received low grades in the extra-class task (see Fig 3 D) than those that evaluated that this task did not contribute to their learning (see Fig. 3 B). This indicates that even those students which received low grades in that task evaluated the task as contributing to their learning, reinforcing that students’ self-evaluation was not driven by their performance.

The increase of students’ self-evaluated learning with students’ perception of teaching effectiveness could be questioned as limited by a biased perception of the students, for example due to affinity or sympathy towards the teacher and towards the teaching methods. However, the increase in the variation of students’ self-evaluated learning with their perception of the effectiveness of teachers’ stimuli (see Fig 3 A), highlights that among the students with high self-evaluated learning, there was a large variation in their perception of teachers’ stimuli. We interpret this as a validation of the detected positive effect of teachers’ stimuli with students’ learning, as far as it shows that students’ evaluation of teachers’ effectiveness does not reflect merely their satisfaction with their learning. Thus, the prejudice stating that students only evaluate positively teachers when they had good results must be discarded.

Our results show that students’ perception was not as accurate as the perspective of their grades, as highlighted by the contrast between the asymptotic relation between students’ grades and their performance in the extra-class group task, while their perception indicated a linear response. Besides, students’ perception did not detect effects of online writing assessments, which were detected through their grades (see Fig 3 E). We suggest that this active learning task — online writing assessments, is more tedious, and thus, less appreciated by the students, which led to a decrease in their perception of this task as contributing to their learning. Thus, although other studies verified students’ ability to detect learning efficiencies [58, 60, 61]; our study shows that students’ perspective on teaching efficiencies, albeit accurate, produces coarse and partial results. Other studies found that, although students learn more in active-class environments, their perception of learning, while positive, is lower than that of their peers in passive environments [77]. When students experience the increased cognitive effort associated with active learning, they initially take that effort to signify poorer learning, which may explain why students did not perceive the online writing assessments as contributing to their learning.

While several studies demonstrated that active learning techniques enhance students’ learning [20, 39], few studies evaluated students’ perception of active learning [21, 78], and none, to our knowledge, compared these two perspectives. As far as learning involves both the teachers and the students, comparing their perspectives is valuable and informs about the mechanisms involved in learning. Our results highlight that these perspectives are complementary, and none of them is overarching.

### Educational challenges in a colonized country

Our study involves cheap but effective methodologies [1], to tackle large sized classes, of diverse socioeconomic and cultural backgrounds. There is a profound need for more effective schools, especially within resource-poor communities in low- and middle-income countries [17], such as Brazil. Besides, there is a specific need for developing ecological consciousness [79], and critical thinking, as a vaccine against fake news and mind manipulation [80]. Here, we applied a strict hypothetical-deductive approach to evaluate explicit hypotheses on the efficiency of active learning tasks and teachers’ behavior, on students’ learning. Our results showed that the perspective of the students and their grades agree, evidencing that active learning tasks and teachers’ stimuli for students’ active involvement enhance students’ learning. We are convinced that active approaches, including the appreciation of students’ individualities, are essential to foster critical thinking, cooperative behavior, tolerance and opening to divergent point of views. Ecology is a discipline that connects areas of knowledge and aims for generalizations with both biological and applied implications, as well as political and economics ones. Therefore, we see as mandatory to go beyond content-centered and teacher-centered learning. Especially in these times of rising pluralities (see Educational challenges in a colonized country for more details), contrasted to a rising intolerance, prejudice, right radicalism, dismiss of environmental issues, and disregard of science itself, as emblematically exemplified by the current Brazilian radical rightist government [66–68], we need a pluralistic, respectful, and student-centred education. We propose that active learning tasks, added to teachers’ behavior stimulating the active involvement of the students, besides increasing students’ learning, are an efficient way to oppose these forces. We recommend ecology teachers to implement multiple simultaneous active learning tasks, including extra-class group task and writing assessments, both as classroom group tasks and as individual online tasks, at home. Our results highlighted that the diversity of tasks is complementary, not redundant. Our practices agree with Handelman’s [35] quest for substituting lecture-based for active-learning based science education.

We are convinced that by acting as a catalyst for students’ active involvement, stimulating and valuing students’ participation, and by implementing active learning tasks, the teacher breaks the domination mechanism of cultural capital [31], giving voice to the students themselves, thus valuing their own cultural capital. Cooperation, as opposed to individuality, is not only a complementary tool for breaking cultural barriers and valuing the voice of the students, but also extends this breakage further by withdrawing the teacher from most of the learning process altogether, thus lessening the burden on the teachers. Active learning tasks stimulate cooperation in the construction of collective knowledge, breaking down barriers for students who do not understand the content, and breaking with the logic that when a student misses or does not know certain content they become inferior to those who are more knowledgeable. Cooperative learning fosters informal peer instruction (see [81, 82] for its formal implementation), as far as the students’ colleague can resolve doubts much more efficiently than the teacher, because of the proximity in language, common cultural universe and similar knowledge level. Active involvement encourages students to discuss questions in groups during a lecture; it breaks with the logic of an authoritarian transmission of knowledge because it makes the student autonomous regarding their learning process. The carrying out of this research is by itself the embodiment of Paulo Freire’s constructivist teaching process of “action-reflection-action” [83–85], as well as Dewey’s reflexive teacher [86–88], as far as teachers, themselves (CFS, RBFC and NS are acting ecology teachers), evaluate their practice and reformulate it upon their observations. For example, CFS based reformulations of the Basic Ecology course on the results of this work.

## Conclusions

Our study shows that students perceive most part of the effectiveness of active-learning tasks and of teachers’ efforts. We showed that different active learning tasks are complementary, so that implementing multiple tasks, together with teachers’ stimuli for students’ active involvement, increase students’ learning. We propose that the diversity of active-learning tasks is important to touch students with differing learning mechanisms, therefore it is a tool for democratizing learning. Based on our results, we recommend that teachers in both college and high school, should implement simultaneous active-learning tasks that include extra-class work in group, individual and group writing assessments, and should stimulate students’ engagement through respectful and non-authoritarian behavior. Furthermore, our results showed that previous basic knowledge also plays a central role in driving effective learning, evidencing the importance of students instruction outside the university, including high school. The effectiveness of most active learning tasks involve cooperative learning, which fuels a virtual circle among teacher’s behavior and students’ commitment, both through extra-class group tasks, and through student’s effort. We emphasize that the applied teaching methodology is cheap and feasible for large classes, and that it is specially important as a tool to foster critical thinking, tolerance and respect for differing points of view. In these times of rising intolerance, prejudice, dismiss of environmental issues, and disregard of science itself, we need an effective, pluralistic, respectful, and student-centred education.

## Supporting information

S1_Text_Survey

S2_Text_Exam1

S3_Text_Exam2

S1_Dataset_Grades

S2_Dataset_Survey

## Acknowledgments

To my colleague teachers in the Basic Ecology (BIO 131) course, José Henrique Schoereder, Flávia Maria da Silva Carmo and Ricardo Ildefonso de Campos (Departamento de Biologia Geral, Universidade Federal de Viçosa), and to the undergraduate and graduate tutors at the course; to the 176 students that “suffered” this experiment; to the ¿ 9000 students that have been enduring our classes in the last 25 years; to the undergraduate Dean of the Universidade Federal de Viçosa, Frederico José Vieira Passos, for the support and for Richard Mark Felder (North Carolina State University) and, Og Francisco de Souza (Departamento de Entomologia, Universidade Federal de Viçosa), for inspiration and valuable suggestions.

## Supporting information

**S1 Text. Anonymous survey on teaching efficiencies.** Google^®^ forms, made available for students one month before the end of the academic semester.

**S2 Text. First exam.** Applied on the 10^th^ teaching week, simultaneously, to all students.

**S3 Text. Second exam.** Applied on the 17th teaching week, simultaneously, to all students.

**S1 Dataset. Students’ self-evaluation.** Answer values of the students for the anonymous survey on teaching efficiencies.

**S2 Dataset. Students’ actual grades.** In the exams and active learning tasks, as well as their effort in the tasks.

## Footnotes

1 In Portuguese we do not distinguish the words “teacher”, for high school, from “professor”, for university. We preferred to maintain all references as “teacher”, so as to emphasize the shared challenges and roles, irrespective of the level of teaching.

2 Grades in the National High-School Exam (ENEM) where 1: *≤* 600 points in the ENEM, 2: 601 to 640, 3: 641 to 700, 4: *≥* 701

3 The National High School Exam includes questions on natural sciences (physics, biology and chemistry), languages (literature, foreign language and Portuguese, arts, physical education and information technology), human sciences (philosophy, sociology, geography and history), mathematics (algebra and geometry) and a separate assessment on writing.

