## Supplementary material for "Effectiveness of active learning for ecology teaching: the perspective of students vs their grades": S1_Text_Survey

Rate BIO 131 Basic Ecology 2015ii

Evaluate the teachers, tutors and structure of the course BIO 131 - Basic Ecology, with grades from 0 (never, terrible) to 5 (always, excellent), as below. Your answers are anonymous, meaning there is no way to identify respondents.
0: never, terrible; 1: almost never, very weak; 2: unsatisfactory; 3: satisfactory; 4: very good, almost always; 5: always, excellent

1. Which is your sex?
   ( ) Male ( ) Female ( ) Other
2. Which is your age?
   ( ) ≤ 18 ( ) 19 a 24 ( ) ≥ 25
3. Where did you live during the 2015ii class?
   ( ) Parents' house
   ( ) Student house
   ( ) Dormitory on campus
   ( ) Alone
   ( ) With spouse / partner
   ( ) With your child / children
4. Did you study high school in public or private?
   ( ) Private school
   ( ) State Public School
   ( ) Federal public school
   ( ) Agricultural Family School (EFA – *Escola Família Agrícola*)
   ( ) Other: ________
5. What is your home State?
6. In which area of knowledge is your undergraduate course inserted?
   ( ) Exact
   ( ) Biological
   ( ) Agrarian
   ( ) Human
7. What was your grade in the National High School Examination (ENEM – *Exame Nacional do Ensino Médio*)?
   ( ) ≤ 600
   ( ) 601 a 640
   ( ) 641 a 700
   ( ) ≥ 701
8. What is your main area of interest at the moment (one word)?
   ________________
9. Have you studied regularly and in advance the content presented? (0 to 5)
10. Have you done the requested activities on time? (0 to 5)
11. Did you have the basic training necessary to achieve good results in the course? (0 to 5)
12. Have you tried to establish a relationship between the content covered and other

contents or facts already known? (0 to 5)

13. Have you tried to interact with the teacher, the tutors and colleagues? (0 to 5)
14. Were you motivated with course? (0 to 5)
15. What is your most frequent way of accessing the internet?
    ( ) Computer laboratories at UFV
    ( ) Smartphone
    ( ) At home
    ( ) Academic internship
    ( ) Others:_______
16. Were you satisfied with your achievement in the course? (0 to 5)
17. Have teachers encouraged questions? (0 to 5)
18. Have teachers seeked active involvement of students beyond simple questions? (0 to 5)
19. Have teachers payed careful attention to student comments, questions and answers and respond constructively? (0 to 5)
20. Have teachers checked periodically if the students are understanding? (0 to 5)
21. Has the extra-class tasks added practical knowledge of the subject matter of theoretical classes? (0 to 5)
22. Do you think that in-class active learning exercises promote the integration of students with more difficulties? (0 to 5)
23. Do you consider that doing the extra-class group task contributed to your learning? (0 to 5)
24. Do you consider that doing the extra-class group task has promoted student cooperation? (0 to 5)
25. Do you consider that  doing the extra-class group task has promoted additional learning domains beyond the classes? Check which.
    ( ) Did not promote any additional learning
    ( ) Increased content learning
    ( ) Worked the affective domain
    ( ) Worked the psychomotor domain (put into practice)
    ( ) Stimulated initiative
    ( ) Stimulated independence
    ( ) Stimulated individual responsibility
    ( ) Stimulated collective responsibility
    ( ) Other: _________
26. Do you consider that redoing exam in pairs contributed to your learning? (0 to 5)
27. Do you consider that online writing assessments contributed to your learning? (0 to 5)
28. Which VIRTUAL PLATFORM OF LEARNING (PVAnet) modules have you enjoyed, have you considered more useful or most interesting?
    ( ) Course description
    ( ) E-mail
    ( ) Reciprocal evaluation of the members of the extra-class group
    ( ) News
    ( ) Assessment corrections and Exam answers

( ) Classroom data-show presentation
( ) Online writing assessments
( ) Extra-class group task steps
( ) Personal presentation
( ) What I learned from the corrections
( ) Doubts
( ) In search of groups / members for the extra-class group task
( ) None
( ) Other:________

29. Which VIRTUAL PLATFORM OF LEARNING (PVAnet) modules did you find most annoying / useless / problematic?
( ) Course description
( ) E-mail
( ) Reciprocal evaluation of the members of the extra-class group
( ) News
( ) Assessment corrections and Exam answers
( ) Classroom data-show presentation
( ) Online writing assessments
( ) Extra-class group task steps
( ) Personal presentation
( ) What I learned from the corrections
( ) Doubts
( ) In search of groups / members for the extra-class group task
( ) None
( ) Other:________
30. Evaluate the utility of the VIRTUAL PLATFORM OF LEARNING (PVAnet) for your learning in this discipline (0 to 5).
31. Do you consider that the objectives of the course have been fully achieved? (0 to 5).
32. Do you consider that the content covered in this course is important for your professional training (0 to 5).
33. Regarding the content and classes of this course, you considered that:
( ) The sequence aroused my interest and showed the applications of theoretical concepts
( ) The classes of the applied part were confusing, because they lacked preliminary theoretical base
( ) There was no closure of applications, gathering all ecological theory
( ) The content is excessive, in relation to the number of classroom time of this course
( ) Classroom lectures were superficial
( ) The mismatch between the order of the classes and the sequence of the textbook made it difficult for me to learn
( ) The classes were dispensable, because I prefer to study everything on my own.
( ) The textbook helped to complement and sediment the concepts seen in class
( ) Other: _______
34. I was informed of the objectives of this research in a clear and detailed manner, and

clarified my doubts. I know that at any time I may request new information and modify my decision to participate if I so desire. I declare that I agree to participate. I received an original copy of this free and informed consent form and was given the opportunity to read and clarify my doubts. The full text of the Free and Informed Consent Form (TCLE) is available at the
<https://drive.google.com/file/d/0B79gNw2VBFgiQ2sxV3dJMHhsY0k/view?usp=sharing>

( ) Yes
( ) No
