## Supplementary material for "Effectiveness of active learning for ecology teaching: the perspective of students vs their grades": S2_Text_Exam1

A

**UNIVERSIDADE FEDERAL DE VIÇOSA**
**CENTRO DE CIÊNCIAS BIOLÓGICAS E DA SAÚDE**
**DEPARTAMENTO DE BIOLOGIA GERAL**
**Prova I de BIO 131 – Ecologia Básica 2015ii**
**09/10/2015 – 8h00 às 10h00**

Nome:_____________________________________________________________ Matrícula: ___________ Turma: _____

**INSTRUÇÕES:**
A prova vale 30 pontos, e é constituída por 15 questões, com 5 afirmativas em cada questão, que você deve responder preenchendo as lacunas com V (verdadeiro) ou F (falso). Assim, cada resposta correta vale 0,4 pontos. As questões 14 e 15 têm 4 itens, cada um valendo 0,5 pontos. Após responder as questões, transfira as respostas para o gabarito, **utilizando uma caneta.**

**BOA PROVA!**

GABARITO

| Número da questão | Respostas | | | | |
|---|---|---|---|---|---|
| 1 | | | | | |
| 2 | | | | | |
| 3 | | | | | |
| 4 | | | | | |
| 5 | | | | | |
| 6 | | | | | |
| 7 | | | | | |
| 8 | | | | | |
| 9 | | | | | |
| 10 | | | | | |
| 11 | | | | | |
| 12 | | | | | |
| 13 | | | | | |
| 14 | ------------ | ------------ | ------------ | ------------ | ------------ |
| 15 | ------------ | ------------ | ------------ | ------------ | ------------ |

**1. A plasticidade fenotípica permite que indivíduos com o mesmo genótipo, sob condições ambientais diferentes, exibam características fenotípicas diferentes. Isso pode acontecer até no mesmo indivíduo. Nas alternativas abaixo, identifique aquelas que se referem corretamente à plasticidade fenotípica, marcando V ou F nas respectivas lacunas**

(   ) Na mesma planta podem existir folhas com espessura e coloração diferentes porque diferentes folhas recebem quantidades diferentes de luz.
(   ) A plasticidade fenotípica é uma propriedade morfo-fisiológica baseada em um conjunto de genes, portanto é selecionável pela seleção natural;
(   ) Mudança sazonal na coloração dos pelos de animais do ártico, como raposas e lebres, que no inverno apresentam coloração mais clara, são exemplos clássicos de plasticidade fenotípica.
(   ) Variações de fenótipos podem ser expressos por um único genótipo sob condições ambientais diferentes, mas sempre dentro de limites geneticamente definidos.
(   ) Mutações também são bons exemplos de plasticidade fenotípica, uma vez que fazem surgir tipos diferentes de indivíduos nas populações.

2. **A ecologia tem estreitas ligações com a evolução e é muitas vezes necessário evocar o pensamento evolutivo para explicar padrões e processos ecológicos. Com base nessa afirmação a respeito da evolução avalie as afirmações abaixo, assinalando V ou F:**

(__) A seleção natural pode ser exercida por fatores bióticos, como a competição e a predação, e também por fatores abióticos do ambiente físico-químico.
(__) As mudanças evolutivas só podem ocorrer através da seleção natural, já que esse fenômeno introduz variação nas populações
(__) As pressões seletivas podem induzir o aparecimento de mutações que conferem maior valor adaptativo para os indivíduos
(__) A mudança da freqüência gênica entre gerações pode ocorrer pela ação de fatores aleatórios que não são seletivos.
(__) A seleção natural privilegia características que evitam que a espécie seja extinta a longo prazo, mesmo que estas características sejam desvantajosas para os indivíduos.

**3. No gráfico abaixo estão representados os limites de tolerância de um organismo à temperatura.**

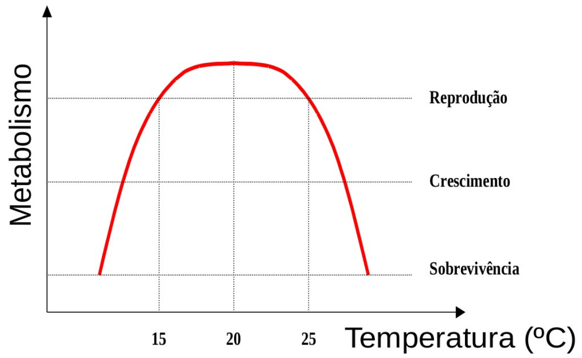


Em relação às informações dispostas no gráfico, preencha as lacunas que precedem as afirmativas com **V** (verdadeiro) ou **F**(falso).

( ) A densidade populacional dessa espécie tende a crescer em locais onde o intervalo de temperatura esteja compreendido entre 15 e 25$^0$C.
( ) Os limites de tolerância à amplitude térmica para esta espécie estão provavelmente entre 10 e 30$^0$C e são parâmetros que definem uma dimensão do nicho da espécie.
( ) Não existem populações dessa espécie em locais com temperatura média abaixo de 15$^0$C, pois os indivíduos não conseguem se reproduzir.
( ) A sobrevivência dos organismos dessa população está condicionada ao aumento da temperatura local para que haja aumento da taxa reprodutiva.
( ) O gráfico mostra que existe uma relação entre o metabolismo dos indivíduos que compõem a espécie e a temperatura local, o que representa uma dimensão do nicho fundamental dessa espécie.

**4. Por que certas aves "gritam" ao perceberem a aproximação de um predador (sinal de alarme)? Relacione as hipóteses explicativas com suas respectivas hipóteses nulas, ou seja, o resultado esperado caso sua hipótese explicativa esteja ERRADA.**

Hipóteses explicativas:
$H_1$: Porque o "grito" assusta o predador.
$H_2$: Porque as aves se assustam com o predador.
$H_3$: Porque essas aves são predadoras do predador.
$H_4$: Porque com isso as aves avisam suas parentes do perigo, aumentando as chances de salvar cópias dos seus genes.
$H_5$: Porque com isso as aves chamam a atenção do predador para si, desviando-o do ninho com seus filhotes.

Responda V para as hipóteses nulas que REFUTEM a hipótese sugerida e F para aquelas que NÃO REFUTEM a hipótese sugerida:

( ) Para H1: Aves sem filhotes também apresentam grito de alerta
( ) Para H2: Aves submetidas a um sinal inesperado não gritam
( ) Para H3: Essas aves incluem sementes em sua dieta
( ) Para H4: O comportamento do predador não é afetado pelos gritos
( ) Para H5: Aves sem filhotes também apresentam grito de alerta

**5. Turchin (2001) propôs uma analogia entre o potencial biótico de crescimento exponencial e a 1ª lei da física mecânica de Newton, a lei da inércia. No raciocínio dessa analogia, seria correto afirmar que (assinale V ou F):**

( ) Enquanto Newton se referia à tendência de movimento de um corpo no espaço, o potencial biótico se refere à tendência de movimento de populações biológicas no espaço.
( ) As populações biológicas se distinguem do universo da física mecânica por não dependerem de reprodução para manterem uma continuidade no tempo.
( ) Enquanto populações biológicas estão sujeitas à evolução, e portanto podem se alterar ao longo das gerações, isso não se aplica a objetos físicos não-vivos.
( ) Segundo a lei da inércia, objetos tendem a manter sua velocidade constante, o que corresponde à tendência biológica de troca de indivíduos (migração) em metapopulações.
( ) Embora o potencial biótico seja inerente a todo o ser vivo, nenhuma população pode manter um crescimento exponencial positivo indefinidamente.

**6. Em termos de dinâmica espacial de populações biológicas, é correto afirmar que (assinale V ou F):**

(   ) Dispersão se distingue da migração porque embora ambos os processos tratem de deslocamento de organismos no espaço, o primeiro é mais abrupto que o segundo.
(   ) Mesmo organismos que eram tidos como de locomoção passiva, como o plâncton, apresentam um processo de migração vertical diário.
(   ) Há três padrões básicos de distribuição espacial – regular, agregado e aleatório – que não dependem da escala espacial da observação.
(   ) Quando uma população local apresenta emigração positiva, ela pode servir de fonte de organismos para outras populações locais.
(   ) O conjunto de populações locais que apresentam uma dinâmica espacial de migração entre si, é chamado de metapopulação.

**7. A figura abaixo mostra a transição demográfica ocorrida na Europa, ao longo dos últimos 100 anos. Em relação ao crescimento e demografia da população humana, é correto afirmar que (assinale V – Verdadeiro ou F – Falso):**

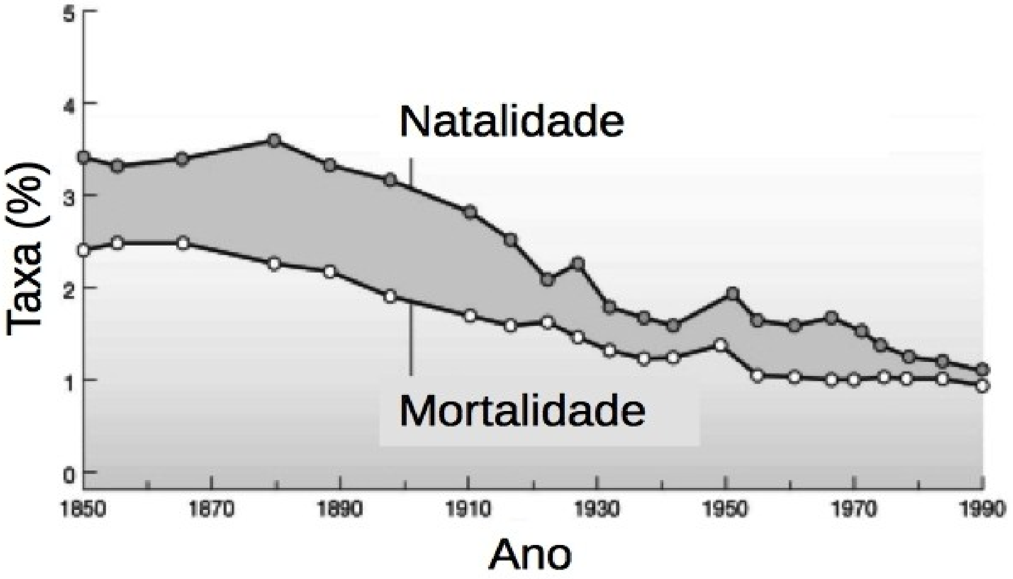


(   ) Nos países mais ricos, tem ocorrido uma transição demográfica, em que a taxa de crescimento populacional tem diminuído
(   ) A transição demográfica corresponde ao aumento nas taxas de natalidade, sendo compensado pela redução na mortalidade
(   ) A diferença entre as duas curvas da figura acima representa a taxa de crescimento populacional
(   ) A expectativa de vida das populações humanas têm aumentado nos últimos 100 anos
(   ) A demografia humana difere muito entre países e regiões do mundo, com maior fertilidade em regiões pobres, e menor fertilidade em países ricos

**8. Em relação à agricultura e sustentabilidade, é correto afirmar que (assinale V – Verdadeiro ou F – Falso):**

(   ) Técnicas de monocultura estão associadas a altos custos com insumos, mecanização e transportes
(   ) Sustentabilidade na agricultura se refere à continuidade de políticas de incentivo fiscal, como redução de impostos, financiamento e linhas de crédito para a exportação
(   ) A teoria ecológica relacionada à sustentabilidade envolve o conceito de estabilidade, resiliência e resistência: quanto maior a resiliência, maior a vulnerabilidade do sistema
(   ) Sistemas agroecológicos necessitam de altos investimentos em maquinário e força de trabalho, de forma a monitorar e reformular técnicas e alternativas
(   ) A produção de alimentos para a redução da fome no mundo depende de investimentos em sistemas produtivos de larga escala

**9. A figura abaixo mostra o colapso dos estoques de bacalhau devido à sobre-exploração. Assim, é correto afirmar que (assinale V – Verdadeiro ou F – Falso):**

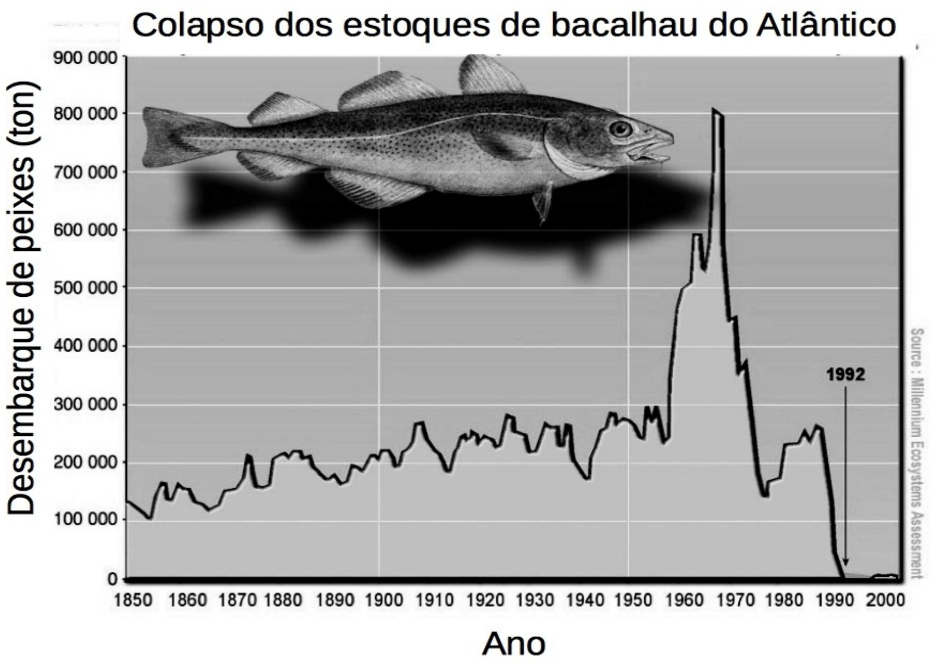


( ) O nível dos estoques pesqueiros no mundo está diretamente correlacionado à intensidade de exploração pela pesca

( ) Para alcançar uma produção máxima sustentada, deve-se retirar a mesma qantidade de peixes que o valor do máximo recrutamento desses organismos

( ) Recrutamento é a diferença entre a natalidade e a mortalidade, sendo máximo quando a taxa de crescimento populacional por indivíduo é máxima

( ) Um dos riscos de colapso ao se utilizar uma estratégia de exploração máxima sustentada é que um alteração ambiental que eleve a capacidade suporte do ambiente pode levar à extinção do recurso

( ) Na exploração por esforço fixo, a quantidade de peixes retirada aumenta com a densidade de peixes

**10. Quanto às relações entre contexto político-econômico e impactos ambientais, é correto afirmar que (assinale V – Verdadeiro ou F – Falso):**

( ) O movimento defendendo a liberalização da sociedade, nas décadas de 1960 e 1970, defendiam bandeiras como "Pelo direito à propriedade privada!" e "Pela redução da carga tributária!"

( ) O colonialismo trouxe o avanço em tecnologias agrícolas para o Brasil e demais países da América Latina

( ) Para que houvesse a expansão das áreas cultivadas com um único produto agrícola, foi fundamental o uso de combustíveis fósseis

( ) A dívida externa dos países subdesenvolvidos resultou em aumento da exportação de matérias-primas e produtos agrícolas

( ) A elevação na qualidade de vida em países ricos implicou em redução na pegada ecológica de suas populações

**11. Poluição pode ser definida como tudo aquilo de que não gostamos quando o ambiente é alterado. Neste sentido, é correto afirmar que (assinale V – Verdadeiro ou F – Falso):**

( ) Ecossistemas sem interferência humana podem sofrer fenômenos de poluição ambiental

( ) A destinação dos cadáveres é uma forma de poluição ambiental que ocorre em colônias de organismos eusociais (abelhas, formigas e cupins)

( ) Os serviços ambientais de detritívoros contribuem para a poluição de pastagens na Austrália

( ) A poluição decorrente de dejetos humanos é controlada com saneamento básico e tratamento de esgotos

( ) A falta de treinamento leva catadores de lixo a aumentarem o impacto ambiental de lixões

**12. A figura abaixo representa os processos de regulação de populações de pragas em quatro regiões agrícolas (i a iv). Com base nessas informações, é correto afirmar que (assinale V – Verdadeiro ou F – Falso):**

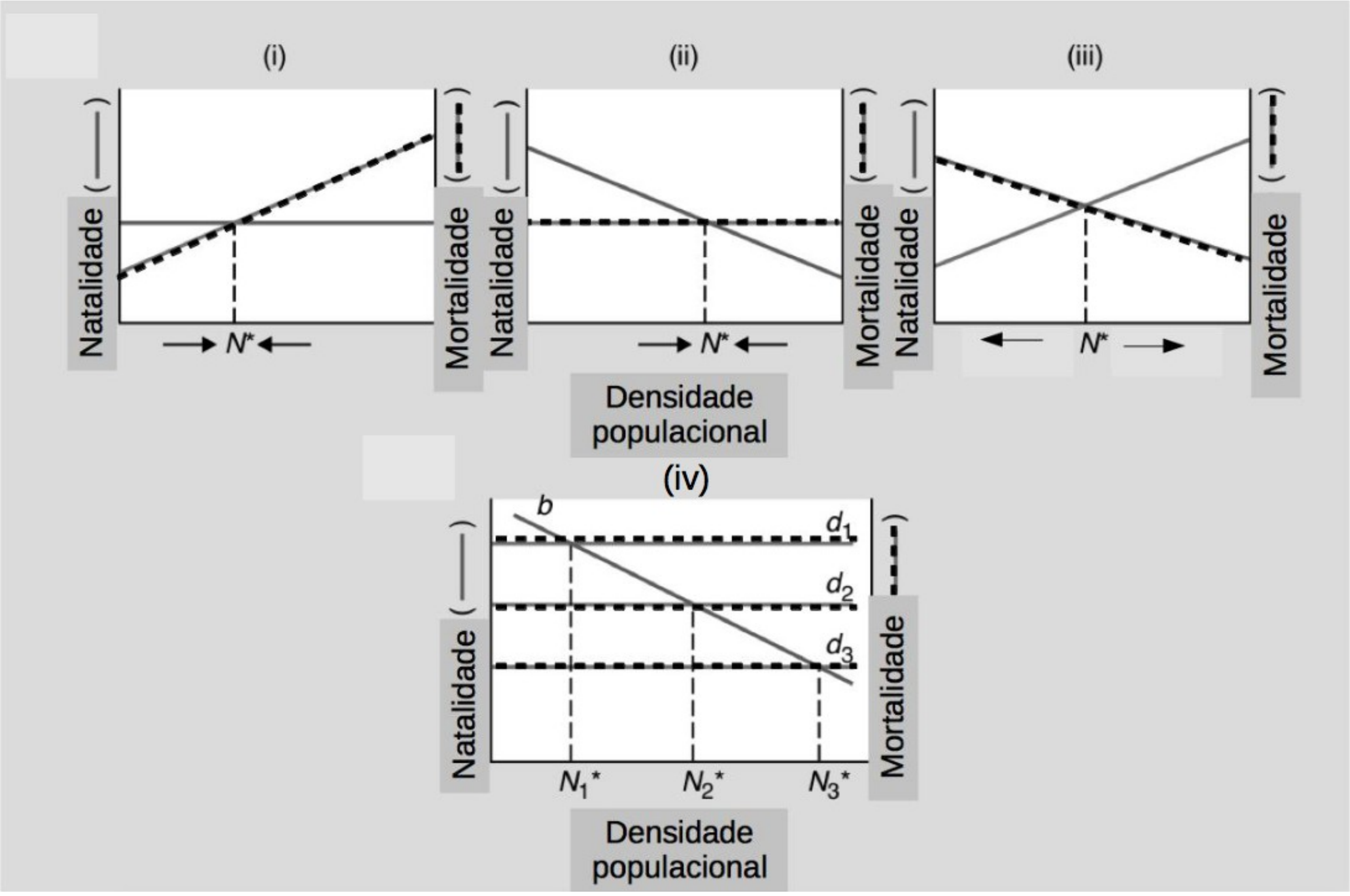


( ) Nas regiões (i), (ii) e (iii) há um equilíbrio estável, em que a população de pragas é mantida na densidade N*, porque as taxas de natalidade são independentes da densidade populacional

( ) Na região (iv), a mortalidade mais alta (d1) leva a uma densidade menor da praga do que a mortalidade mais baixa (d3)

( ) Quando a natalidade diminui com a densidade ou a mortalidade aumenta com a densidade, há uma densidade estável da população

( ) Na região (iii), populações menores que N* podem ir à extinção, porque nessa situação a natalidade é maior que a mortalidade

( ) A figura (iv) mostra como fatores independentes da densidade podem regular populações em diferentes densidades

**13. No último número da revista *Science*, de setembro de 2015, foi publicado trabalho mostrando que nos últimos 40 anos o comprimento da "língua" de abelhas têm diminuído (figura abaixo), provavelmente em resposta à redução na disponibilidade de flores, correlacionada ao aquecimento global. Os autores sugerem que a redução no comprimento da "língua" seja uma forma de reduzir o custo energético para locomoção, um carácter favorecido nos ambientes mais recentes, em que ocorreu redução na densidade de flores. Sobre essas mudanças pode-se dizer (V – Verdadeiro ou F – Falso):**

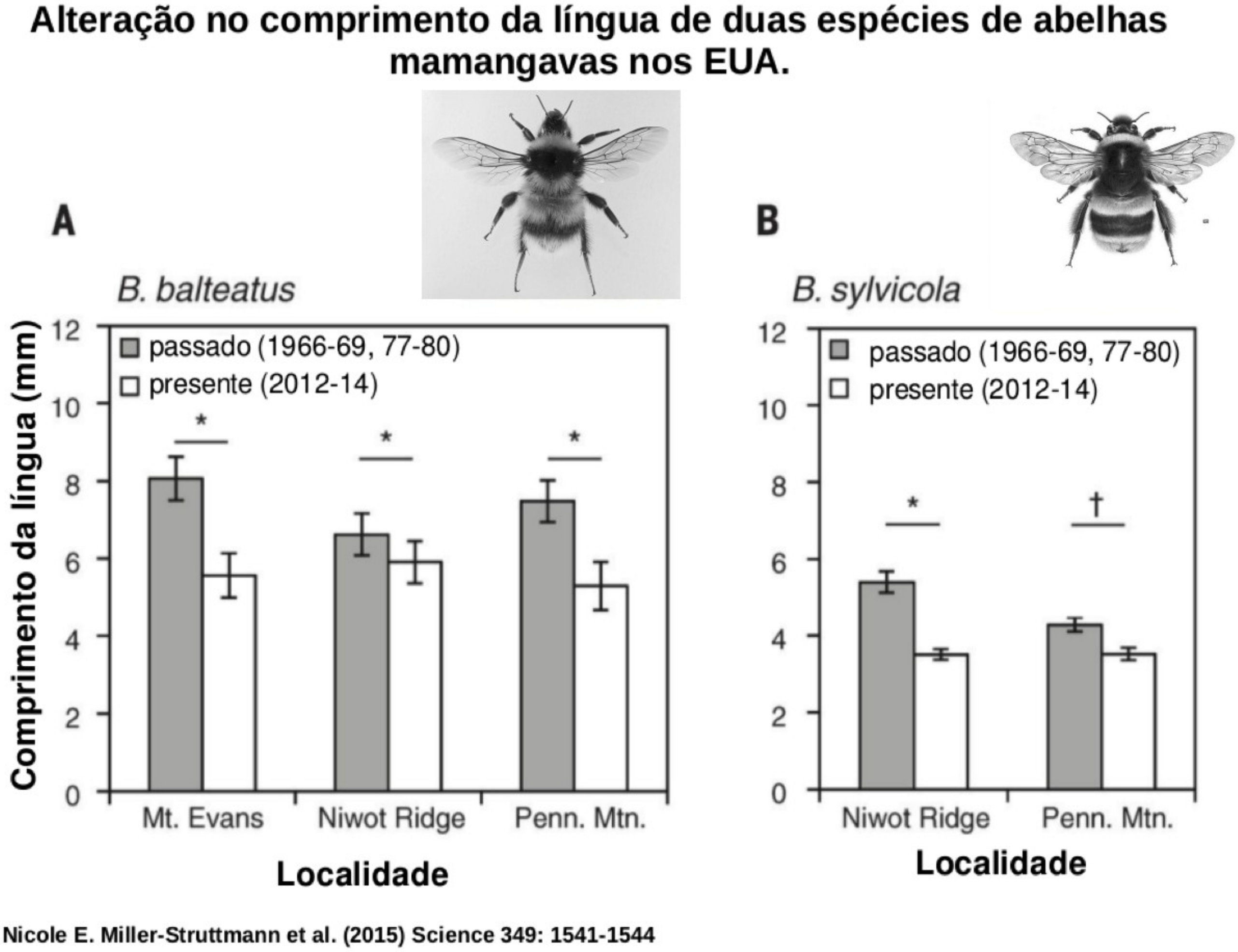


(    ) As duas espécies de abelhas estudadas apresentaram tendências opostas, em relação ao comprimento de suas línguas no "passado" *versus* "presente".

(    )  Para aceitar a hipótese de que a redução no comprimento das línguas ocorreu em resposta à redução na disponibilidade de flores, é necessário mostrar que, no mesmo período, a disponibilidade de flores aumentou.

(    ) Os gráficos acima são um exemplo de variável explicativa categórica, em que as barras verticais mostram a variação dos dados em cada grupo analisado.

(    )  As mudanças observadas nas abelhas podem resultar de flutuação aleatória decorrente de deriva genética, associada à redução das populações de abelhas.

(    ) Uma vez que o comprimento das línguas das abelhas determina a profundidade das flores das quais as abelhas são capazes de retirar néctar, as mudanças observadas nas abelhas podem reduzir o número de abelhas que visitam flores mais profundas.

**14.** Para que uma população esteja regulada, em torno de uma densidade estável, ela precisa apresentar natalidade, mortalidade ou ambos, dependentes da densidade. Complete os gráficos abaixo, **desenhando como deveria ser a variação da taxa de mortalidade com a densidade**, **e como seriam as setas do tempo**, para que (a primeira alternativa (e) é um exemplo):

(e) a população apresente dois pontos de equilíbrio, sendo o segundo um ponto de equilíbrio estável, numa densidade maior, e o primeiro um ponto de equilíbrio instável, abaixo do qual ocorre a extinção desta população

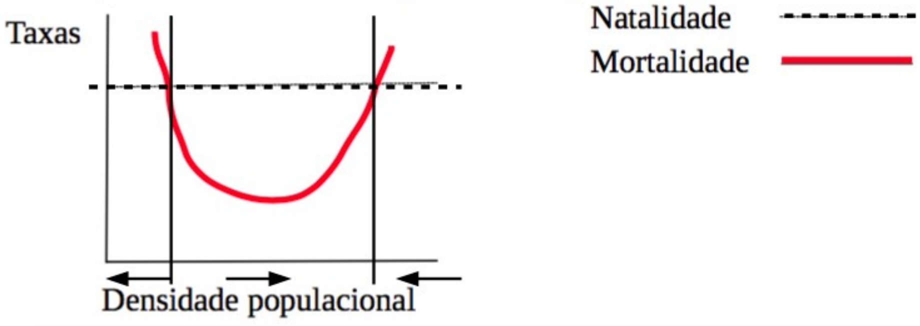


(a) haja um ponto de equilíbrio estável, que corresponde à capacidade suporte (K) da população:

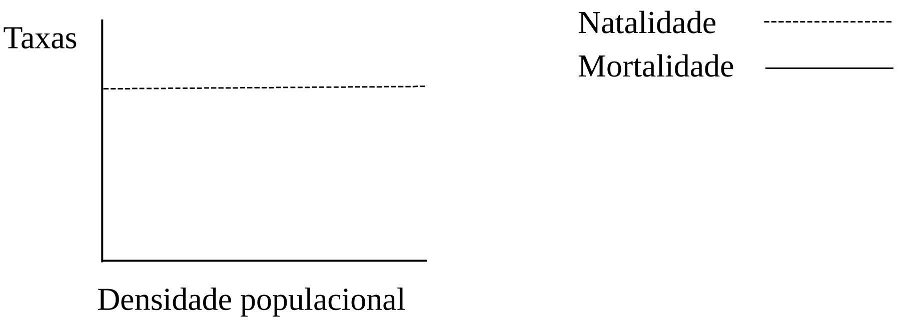


(b) haja uma densidade mínima viável (População mínima viável), abaixo da qual a população irá à extinção:

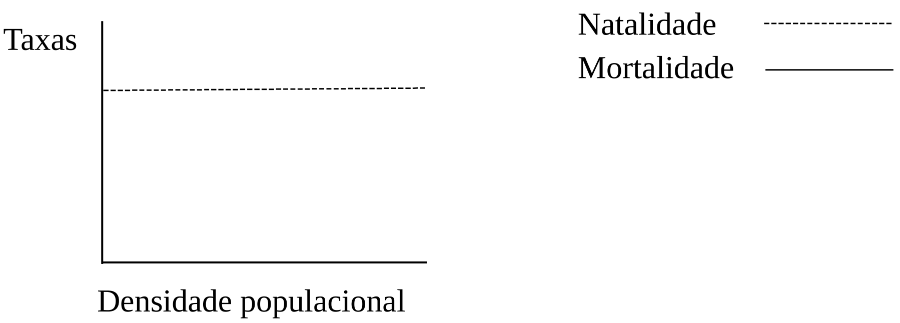


(c) haja uma densidade máxima instável, acima da qual a população apresentará um surto populacional:

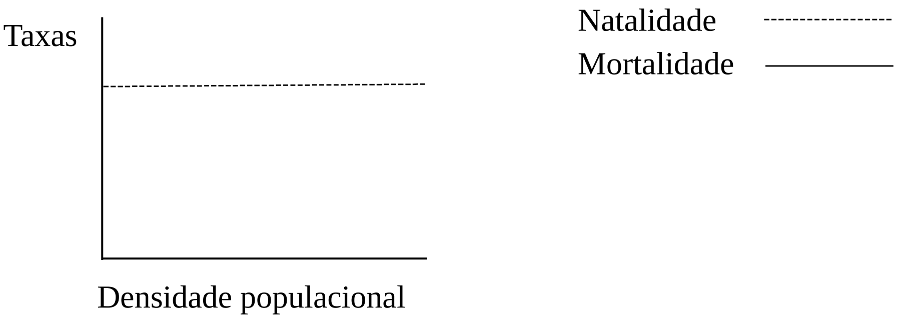

(d) a população apresente dois pontos de equilíbrio, sendo o primeiro ponto numa densidade menor, estável, e o segundo um ponto de equilíbrio instável, com surto populacional acima dele

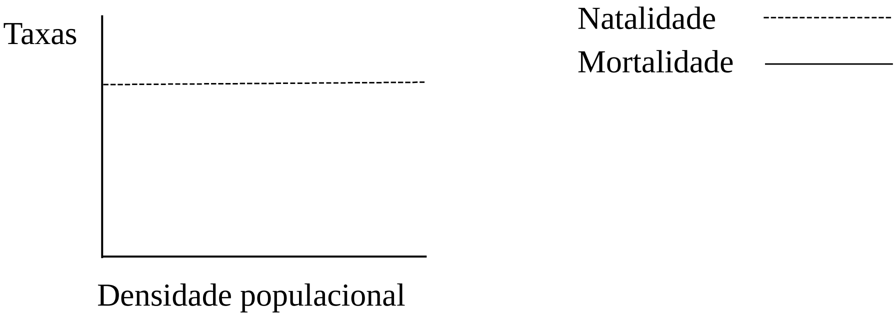


**15. A seguir apresentamos o número de indivíduos e de espécies de grilos em sete fragmentos florestais diferentes da região de Foz do Iguaçu, PR.**

(a) A partir dos dados da tabela ao lado, construa um gráfico da resposta do número de espécies de grilo ao número de indivíduos, plotando os pontos para cada linha da tabela.
(b) Coloque a legenda dos eixos no gráfico.
(c) No mesmo gráfico, trace uma curva esperada para o efeito do número de indivíduos sobre o número de espécies, que obedeça a uma função linear y = a + b.x.

Número de indivíduos e de espécies de grilo em 9 fragmentos florestais na região de Foz do Iguaçu, PR.

| Local | Indivíduos | Espécies |
|---|---|---|
| A | 98 | 41 |
| B | 66 | 31 |
| C | 6 | 28 |
| D | 2 | 25 |
| E | 76 | 1 |
| F | 38 | 46 |
| G | 68 | 4 |
| H | 2 | 36 |

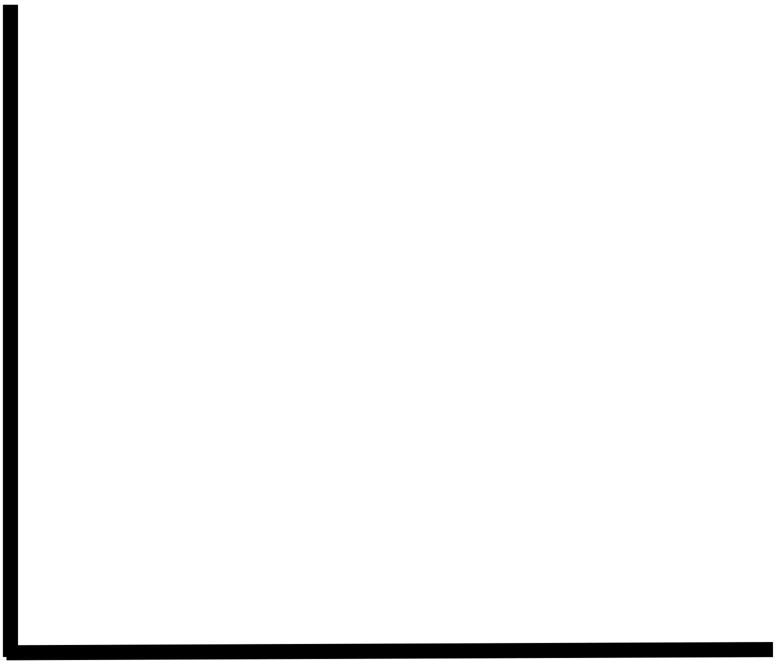

(d) Descreva o efeito observado do número de indivíduos sobre o número de espécies. (máx. 15 palavras)

### FOLHA EM BRANCO
