## Supplementary material for "Effectiveness of active learning for ecology teaching: the perspective of students vs their grades": S3_Text_Exam2

A

**UNIVERSIDADE FEDERAL DE VIÇOSA**
**CENTRO DE CIÊNCIAS BIOLÓGICAS E DA SAÚDE**
**DEPARTAMENTO DE BIOLOGIA GERAL**
**Prova II de BIO 131 – Ecologia Básica 2015ii**
**27/11/2015 – 8h00 às 10h00**

Nome:__________________________________________ Matrícula: ___________ Turma: _____

**INSTRUÇÕES:**

A prova vale 30 pontos, e é constituída por 12 questões, com 5 afirmativas em cada questão, que você deve responder preenchendo as lacunas com V (verdadeiro) ou F (falso). Assim, cada resposta correta vale 0,5 pontos. Os dois primeiros itens da **questão 12** devem ser respondidos **na última folha**. Após responder as questões, transfira as respostas V ou F para o gabarito, **utilizando uma caneta.**

**BOA PROVA!**

GABARITO

| Número da questão | Respostas | |
|---|---|---|
| 1 | | |
| 2 | | |
| 3 | | |
| 4 | | |
| 5 | | |
| 6 | | |
| 7 | | |
| 8 | | |
| 9 | | |
| 10 | | |
| 11 | | |
| 12 | ------ | ----- |

**Questão 1. "**Há preocupação crescente de que a degradação ambiental acelerada ameace interações mutualísticas. Como mutualismo pode conectar espécies a um destino comum, a quebra de mutualismos tem o potencial de expandir e acelerar os efeitos de mudanças globais sobre a perda de biodiversidade e o rompimento de ecossistemas (Kiers e colaboradores (2015) *Ecology Letters 13*: 1459-1474). No contexto de interações mutualísticas e sua trajetória evolutiva, assinale Verdadeiro (V) ou Falso (F):

( ) A redução na abundância de espécies mutualistas chave, como morcegos frugívoros, que agem como dispersores de sementes, pode levar a um efeito cascata, em que as plantas que fornecem os recursos para esses morcegos, podem ser extintas

( ) Relações entre endoparasitas e seus hospedeiros podem se transformar ao longo do tempo evolutivo, com redução da virulência do patógeno e aumento da tolerância do hospedeiro. Essa é a hipótese atualmente aceita para a origem de cloroplastos e mitocôndrias

( ) Um dos efeitos das mudanças climáticas globais é a perda de sincronia entre floração e seus polinizadores, o que pode levar à extinção dos polinizadores devido a escassez de polinização cruzada, consequente redução na variabilidade genética e depressão endogâmica

( ) Mesmo espécies predadoras podem funcionar como espécies do tipo "pedra angular", quando sua ação reduz a densidade de presas competitivamente inferiores, evitando, assim, a exclusão competitiva de espécies competitivamente superiores

( ) Damos o nome de anéis miméticos a conjuntos de espécies de borboleta com coloração parecida, que se beneficiam ou prejudicam entre si em diferentes graus, dependendo de sua frequência em relação às outras, e dependendo do seu grau de toxicidade

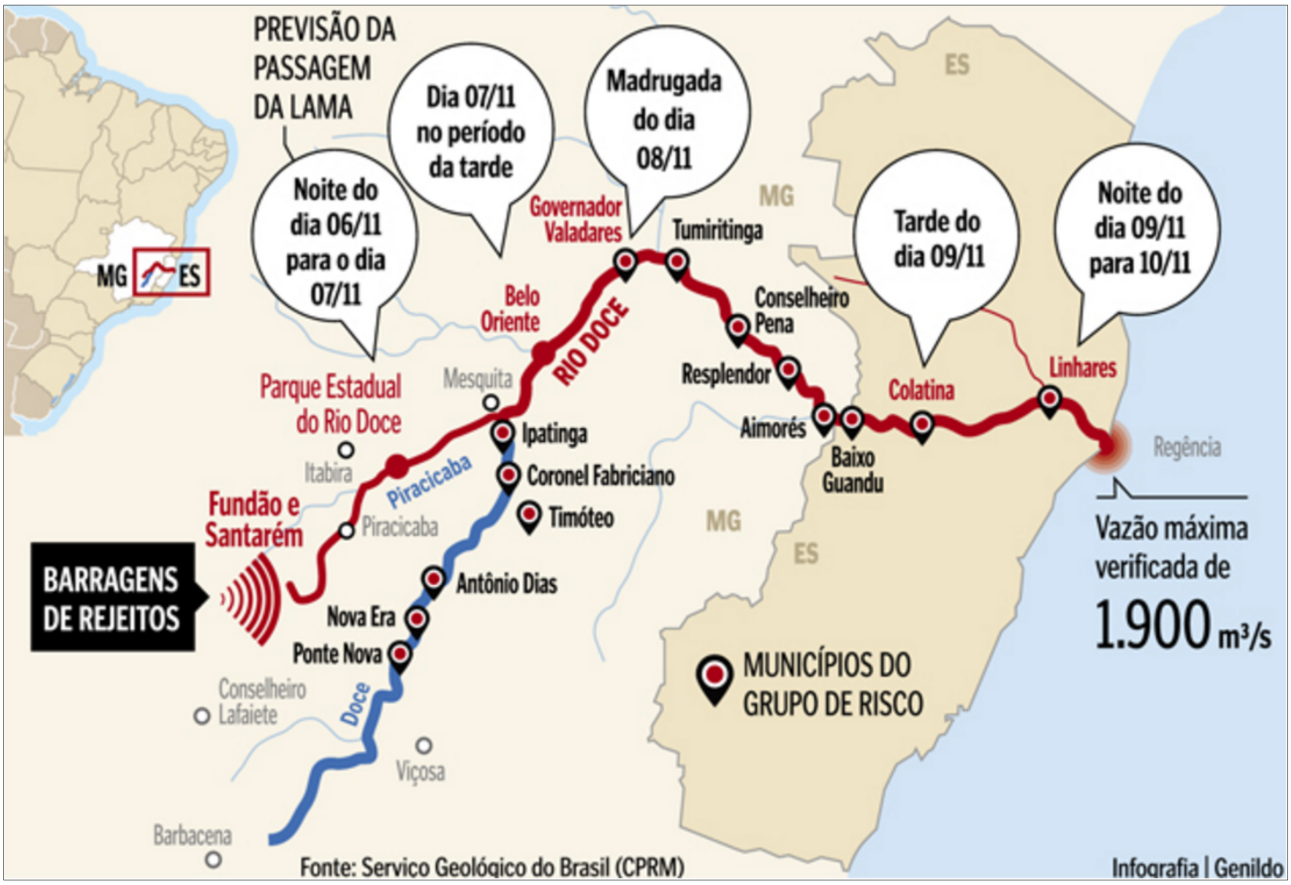

**Questão 2.** A Figura acima representa parte da Bacia Hidrográfica do Rio Doce, que inclui as cidades de Mariana, Ponte Nova e Viçosa. A partir das informações nessa figura e do que você aprendeu sobre o Desastre Ambiental de Mariana, é correto afirmar que (assinale V – verdadeiro ou F – falso):

( ) Apesar de se encontrarem dentro da bacia do Rio Doce, as cidades de Ponte Nova e Viçosa não foram alcançadas pela lama da barragem de rejeitos porque se localizam a montante (rio acima)

( ) Os impactos diretos do desastre se estendem a toda a bacia hidrográfica do Rio Doce, pois os rejeitos estão sendo carreados pelos rios

( ) Os impactos indiretos do desastre envolvem a fragilização econômica e de suprimento de água potável, mas não ameaça o suprimento de água para a indústria e agricultura

( ) As medidas tomadas para a contenção dos impactos incluem a instalação de barreiras filtradoras para a lama e resíduos sólidos carreados pelo rio

( ) O resgate de fauna do Rio Doce, que envolve a captura de peixes dos locais contaminados, e sua liberação em locais “saudáveis”, pode levar à extrapolação da capacidade suporte do local destino, levando à exclusão competitiva de organismos introduzidos ou nativos

**Questão 3.** Vimos que há dois modelos fundamentais de crescimento populacional, um para crescimento ilimitado, o modelo exponencial, e um modelo para crescimento limitado, o modelo logístico. Vimos que ambos são simplificações da natureza, mas que nos permitem compreender mecanismos populacionais, assim como modificar os modelos para aproximá-los do que observamos na natureza. Abaixo apresentamos os dois modelos e suas previsões para o crescimento populacional. É correto afirmar que (assinale V – verdadeiro, ou F – falso):

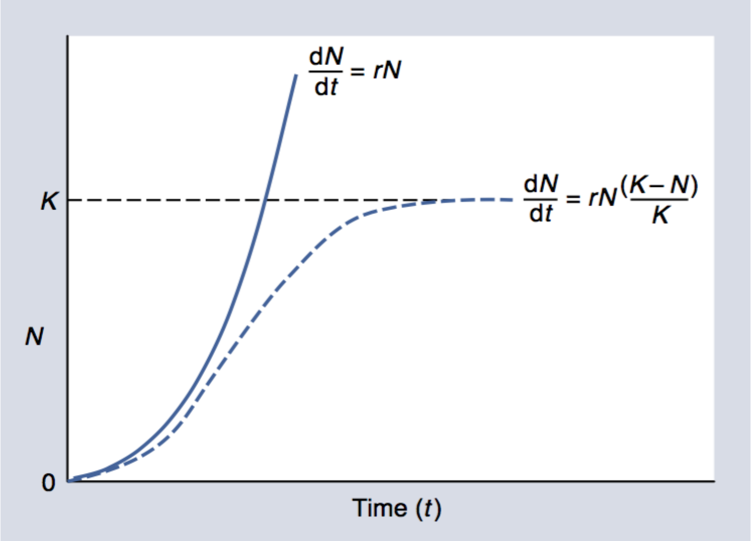

( ) O modelo exponencial representa o potencial biótico, em que toda população biológica cresce em progressão geométrica, quando sua taxa de reposição individual (taxa de reprodutiva *per capita*, $\lambda$ ) é maior do que um

( ) No modelo exponencial, caso a natalidade for menor que a mortalidade, a população diminui até sua extinção

( ) A taxa de crescimento populacional *per capita*, $\frac{dN}{N.dt}$, é uma função linear da densidade populacional nos dois modelos

( ) No modelo logístico, quando N > K, a taxa de crescimento populacional é igual a zero

( ) Quando se introduz retardo temporal no modelo logístico, ao invés de uma estabilização populacional quando N = K, o modelo prevê uma flutuação atenuada, com a população ultrapassando K e sendo reduzida abaixo de K alternadamente

**Questão 4.** Um levantamento de plantas realizado em áreas de campo com tempo de sucessão autotrófica variando de 1 a 40 anos, gerou os resultados sumarizados nas curvas de abundância da figura abaixo. Símbolos vazios são herbáceas, semi-preenchidos são arbustos e totalmente preenchidos são árvores. Acerca desse gráfico responda:

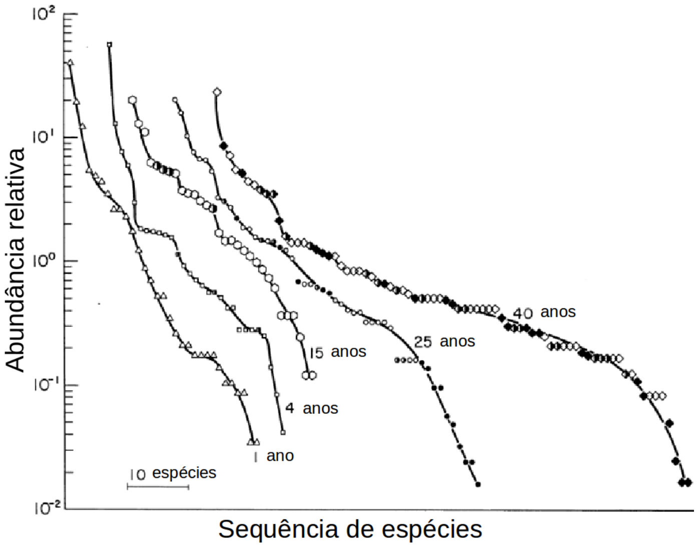

( ) Houve um aumento na riqueza de espécies com a sucessão

( ) A área com com maior dominância tem 40 anos de idade

( ) A figura mostra que a composição de espécies se alterou ao longo da sucessão

( ) A figura não permite avaliar a resposta da riqueza de espécies à sucessão

( ) As espécies arbóreas foram sendo substituídas por herbáceas, ao longo da sucessão

**Questão 5.** A infecção pelo vírus **ebola** causa uma **febre hemorrágica**, uma das doenças virais mais perigosas, frequentemente fatal, com índice de mortalidade de 50 a 90% dos casos. O patógeno é transmitido pelo contato direto com o sangue, secreções ou sêmen de pessoas portadoras do vírus. Frequentemente, funcionários da saúde que mantém contato direto com doentes ou mortos, são infectados. Pelo sêmen a transmissão pode ocorrer até sete semanas após a recuperação clínica da doença. As populações africanas são infectadas em alto número, devido à cultura das aldeias, em que as famílias tem o costume de lavar o corpo dos mortos de forma manual e com cuidado antes do enterro. Assim, o indivíduo morto pelo ebola, transmite o vírus a todos aqueles que tiverem contato com o seu corpo. A profilaxia compreende o isolamento total dos infectados. O vírus ebola, considerado por muitos, o vírus mais perigoso que a humanidade conhece, é um filovírus de forma filamentosa que não possui classificação. Ele recebeu essa denominação porque foi identificado pela primeira vez em 1976 na República Democrática do Kongo (antigo Zaire), perto do Rio Ebola. Sabe-se, atualmente que, o vírus ebola não é altamente infeccioso. Por isso, é muito difícil ocorrer uma epidemia nos países ocidentais, pois a higiene bloqueia qualquer expansão de casos de transmissão do vírus de uma pessoa para outra. **Com base nestas informações, e em seus conhecimentos sobre evolução de interações biológicas, responda se as afirmativas abaixo podem ser verdadeiras (V) ou falsas (F).**

( ) Para que haja evolução de cepas menos virulentas do patógeno, é necessário que haja variabilidade genética relacionada ao grau de virulência do vírus, ou que ocorra uma mutação que codifique menor virulência no vírus
( ) A seleção natural favorecerá cepas do vírus que reduzam a mortalidade dos hospedeiros, de forma a promover a coexistência harmoniosa das duas espécies
( ) Medidas profiláticas de higiene e isolamento de hospedeiros infectados poderão exercer pressão seletiva sobre o vírus, favorecendo cepas mais resistentes à doença
( ) Uma medida profilática eficiente seria a seleção de hospedeiros resistentes ao vírus
( ) Em aldeias com alta proporção de infectados pelo vírus ebola, pequena redução da virulência em algumas cepas do vírus podem ter alto valor adaptativo para o vírus

**Questão 6.** A figura abaixo mostra a distribuição de 4 espécies de roedores escavadores de solo, em diferentes profundidades de um mesmo ambiente. A partir dessa figura, assinale verdadeiro – V ou falso – F:

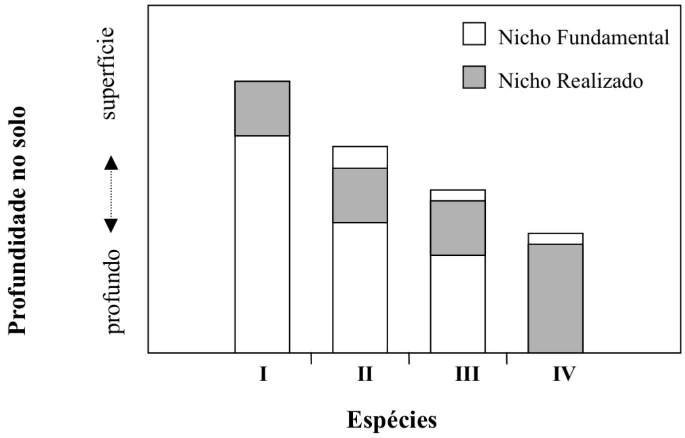

( ) I tem o limite de tolerância mais amplo e IV é a espécie competitivamente superior dentro de seus limites de tolerância.
( ) I, II e III têm os mesmos limites de tolerância e IV é competitivamente superior dentro de seus limites de tolerância.
( ) I é competitivamente superior dentro de seus limites de tolerância e IV tem os limites de tolerância mais amplos.
( ) I tem o limite de tolerância mais amplo e é competitivamente superior dentro de seus limites de tolerância.
( ) III tem os limites de tolerância mais amplos e IV é competitivamente superior dentro de seus limites de tolerância.

**Questão 7.** Abaixo estão listados os principais tipos dinâmica espacial de populações biológicas. Em relação a isso, assinale V – verdadeiro ou F – falso:

A. Predomínio de uma única população perene, que não se extingue, e que fornece colonizadores que resgatam populações menores, essas sujeitas a extinção frequente.
B. Populações locais interligadas por frequentes trocas de indivíduos, nas quais as populações locais nunca sofrem extinção.
C. Populações locais que frequentemente sofrem extinção, e são resgatadas por imigração de indivíduos de outras populações locais.
D. Populações locais com natalidade superior à mortalidade, que fornecem colonizadores que resgatam populações locais com mortalidade superior à natalidade.
E. Populações locais com frequentes extinções, em que a troca de indivíduos é menor que a frequência de extinções, de forma que o resgate é insuficiente para compensar as extinções locais.

( ) Metapopulações clássicas são todas as populações de populações em que a extinção é maior que o resgate populacional
( ) As letras maiúsculas acima correspondem aos seguintes tipos de metapopulações: A. Clássica, B. Manchada, C. Fonte-sumidouro, D. Resgate, E. Não equilíbrio
( ) A degradação e fragmentação de hábitat podem transformar populações B em A, C, D e E
( ) Em termos de conservação biológica, ao se detectar populações com dinâmica espacial como em C, deve-se priorizar a manutenção da conectividade entre populações locais
( ) Caso se trate de espécies prejudiciais ao ser humano, uma dinâmica espacial na forma descrita em E é a mais prejudicial, pois implica em risco permanente de ressurgência

**Questão 8.** Por que alguns locais têm mais espécies do que outros? Em relação à coocorrência de espécies, é correto afirmar que (assinale V ou F):

( ) Um local pode ter maior riqueza de espécies porque tem maior disponibilidade de recursos
( ) Em comunidades saturadas, a competição interespecífica regula a amplitude e sobreposição dos nichos
( ) Caso prevaleçam processos regionais na regulação da riqueza de espécies, haverá um equilíbrio dinâmico entre imigração e extinção
( ) Predadores de topo podem permitir aumento na sobreposição de nichos, por limitarem espécies competitivamente superiores
( ) O aumento da riqueza de espécies com a produtividade é uma evidência favorável à hipótese de que as comunidades biológicas são saturadas

**Questão 9.** É correto afirmar que (assinale verdadeiro – V ou falso – F):

( ) O processo evolutivo de origem de espécies envolve, necessariamente, a interrupção do fluxo gênico.
( ) A especiação simpátrica é o processo evolutivo de seleção natural que favorece genótipos que produzam mais filhotes.
( ) A riqueza de espécies aumenta com a redução na área amostrada.
( ) A maior diversidade em regiões tropicais pode ser explicada pela maior quantidade de energia que flui nestes ecossitemas.
( ) Quanto maior a heterogeneidade ambiental, maior o número de espécies.

**Questão 10.** A coocorrência de espécies pode ser explicada por (assinale verdadeiro – V ou falso – F):

( ) Em comunidades saturadas, a competição não regula a coocorrência de espécies.
( ) Se o recurso for limitante, comunidades com espécies mais especializadas suportam mais espécies do que comunidades com espécies mais generalistas.
( ) A coocorrência de espécies competidoras pode ser mediada pela ação de distúrbios ambientais, que mantém as populações acima do nível competitivo.
( ) A coocorrência de espécies é o resultado de processos regionais de imigração e processos locais de extinção.
( ) Em comunidades saturadas, espécies em locais mais ricos tendem a reduzir sua similaridade, utilizando porções distintas de seu nicho potencial.

**Questão 11.** A figura abaixo representa a resposta de três comunidades à mesma perturbação. A perturbação é representada pela seta vertical, e corresponde a um tufão. A partir desta figura, assinale verdadeiro – V ou falso – F:

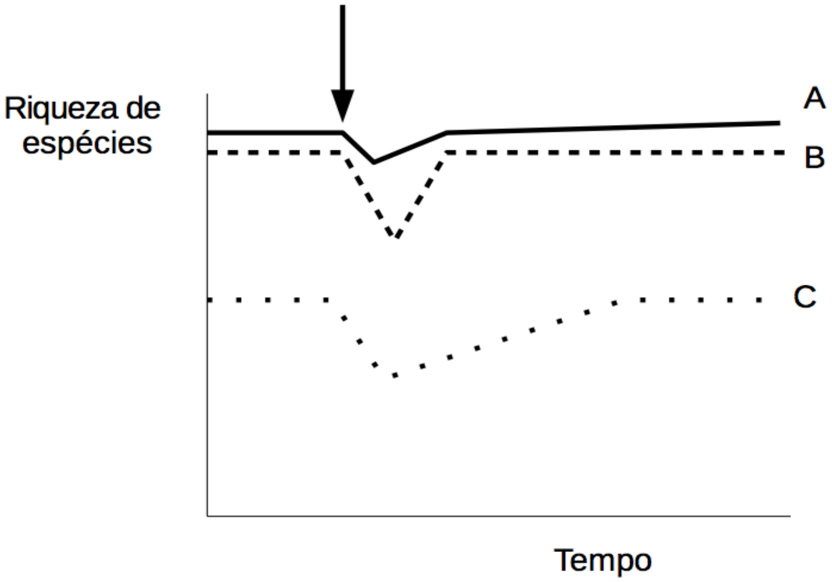

( ) A estabilidade diminuiu com a riqueza de espécies da comunidade.
( ) A comunidade C teve menor elasticidade (=resiliência) que as comunidades A e B.
( ) A comunidade A teve maior resistência do que as comunidades B e C.
( ) As três comunidades foram resilientes à perturbação.
( ) O tempo de recuperação das comunidades foi negativamente correlacionado à sua riqueza de espécies.

**Questão 12.** O mapa abaixo representa um sistema de ilhas próximas da costa de um continente. As ilhas A, B e C estão à mesma distância do continente (e também estão à mesma distância da ilha D). Já a ilha D é a mais distante do continente. As ilhas A e D têm a mesma área, e são maiores do que C, que por sua vez é maior do que B.

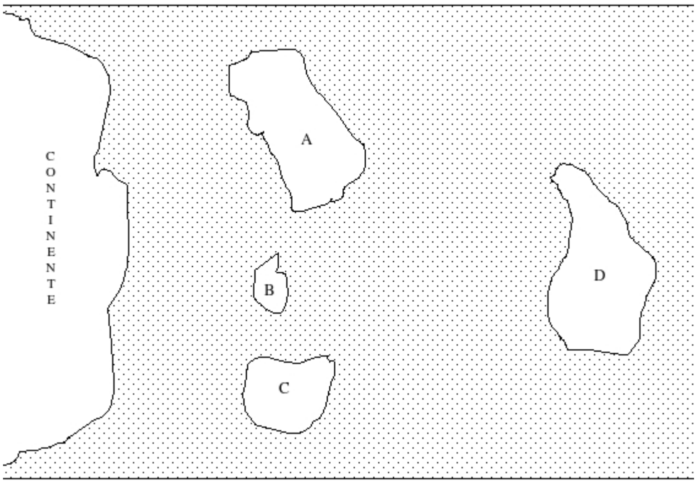

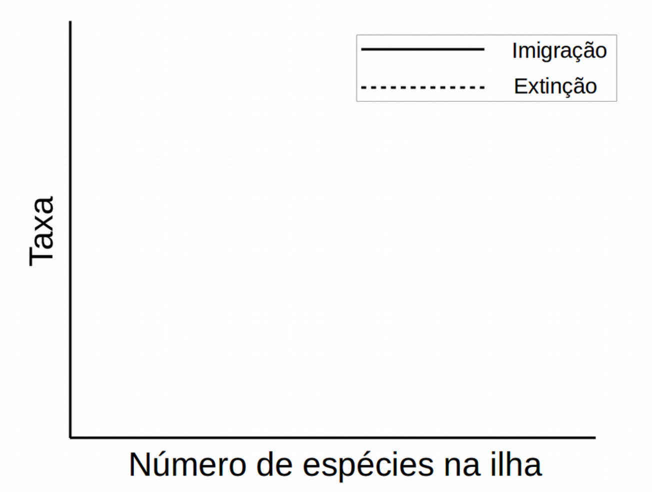

(a) A partir dos dados acima, complete a figura à direita, assinalando as letras das ilhas para cada uma das 8 curvas de extinção e imigração.
(b) Assinale o número de espécies em equilíbrio para cada ilha, no eixo X.

A seguir, assinale verdadeiro (V) ou falso (F):

( ) Na intersecção entre imigração e extinção, ocorre um equilíbrio instável da riqueza de espécies na ilha
( ) A taxa de imigração é mínima quando a riqueza de espécies na ilha for igual à do continente
( ) O efeito resgate representa a interação entre imigração e extinção

FOLHA EM BRANCO

FOLHA EM BRANCO
